# Insights into deuterostome evolution from the biphasic transcriptional programme of hemichordates

**DOI:** 10.1101/2022.06.10.495707

**Authors:** Alberto Perez-Posada, Che-Yi Lin, Tzu-Pei Fan, Ching-Yi Lin, Yi-Chih Chen, José Luis Gómez Skarmeta, Jr-Kai Yu, Yi-Hsien Su, Juan J. Tena

## Abstract

Evolutionary history of deuterostomes remains unsolved and is intimately related to the origin of chordates. Among deuterostomes, hemichordates and echinoderms (collectively called Ambulacraria) are sister groups of chordates. Comparative studies involving these three groups provide valuable insights into deuterostome evolution. Indirect developing hemichordates produce planktonic larvae that bear resemblance to echinoderm larvae before undergoing metamorphosis into an adult body plan with anteroposterior polarity homologous to that of chordates. Therefore, understanding the developmental processes of indirect-developing hemichordates can help understand the evolution of deuterostomes and the origins of chordates. In this study, we analysed the transcriptomes and chromatin accessibility of multiple developmental stages of the indirect-developing hemichordate *Ptychodera flava* and discovered that it exhibits a biphasic developmental program controlled by distinct sets of transcription factors and their corresponding regulatory elements. Comparative analyses of transcriptomes and network analyses revealed that the gastrula transcriptome is relatively ancient, and the TFs orchestrating its gene expression are highly interconnected in networks of cis-regulatory interactions. Comparing the developmental transcriptomes of hemichordates, echinoderms, and amphioxus, revealed high conservation of gene expression during gastrulation that extends to the neurula stages of amphioxus, along with remarkable similarity in larval transcriptomes across the three species. Additionally, we show that *P. flava* possesses conserved interactions of transcription factors necessary for the development of echinoderm endomesoderm and chordate axial mesoderm, including conserved cis-regulatory elements of the FoxA transcription factor that is central to the two networks. These findings suggest the existence of a deuterostome phylotypic stage during gastrulation governed by gene regulatory networks with conserved cis-regulatory interactions. Conversely, integration of gene expression data with synteny data revealed that gene expression recapitulates the independent evolutionary history of the Ancestral Linkage Groups that underwent rearrangements in each deuterostome lineage, suggesting a potential role of genome rearrangement during the evolution of larval strategies in hemichordates and deuterostome body plans.

## Introduction

Deuterostomes comprise a major group of animals that share several unique morphological and developmental features, including radial cleavage, deuterostomy, enterocoely formation of the mesoderm, mesoderm-derived skeletal tissues, and the presence of pharyngeal openings or slits (Satoh 2016; Gee 2018; Nanglu et al. 2023). Despite monophyly recently challenged by phylogenomic studies, current consensus still considers Deuterostome lineages diverged very early after the emergence of bilaterians, and thus may have retained many ancestral traits inherited from the Bilaterian common ancestor (Simakov et al. 2022; Kapli et al. 2021). Therefore, knowledge of the genetic controls underlying the development of representative deuterostome models holds the potential to provide insights into the early origins of both deuterostome and bilaterian animals. Comparative studies of species at key positions of the deuterostome tree could additionally help to understand the complex evolution of these animals, and contribute to reveal genetic and developmental mechanisms underlying the process of diversification and the emergence of evolutionary novelties.

Within deuterostomes, hemichordates and echinoderms are grouped in Ambulacraria based on developmental and molecular data, thus representing the sister group to chordates (Halanych et al. 1995; Cameron, Garey, and Swalla 2000; Eric Röttinger and Lowe 2012). Despite morphological differences of the adult body plan, several hemichordate groups undergo indirect development with free-swimming, planktonic larvae like those of most echinoderms (Lowe 2021; McClay 2011). Similar patterning mechanisms, complemented with conserved expression patterns of key regulatory genes, have been shown to control the formation of the dorsoventral (DV) axis, mesoderm, nervous system and primordial germ cells in echinoderm and hemichordate larvae (E. Röttinger and Martindale 2011; Eric Röttinger et al. 2015; Chang et al. 2016; Fan et al. 2018; Lin, Yu, and Su 2021; Su et al. 2019). This poses the question of whether indirect development was a trait present in the common ancestor of both animal lineages. On the other hand, in contrast to pentaradial symmetry in adult echinoderms, hemichordates share bilateral symmetry with chordates, with a distinctive tripartite body plan of proboscis, collar and trunk (Urata and Yamaguchi 2004; Billie J. Swalla 2007). They show conserved structures such as pharyngeal gill slits, supporting their position within deuterostomes (Simakov et al. 2015; Lowe et al. 2015). Importantly, numerous hemichordate structures exhibit conserved patterning mechanisms during development. These include the overall anteroposterior (AP) axis regionalisation (Lowe et al. 2003) and the AP patterning of the ectoderm in the proboscis and collar by signalling organising centres similar to those of the vertebrate central nervous system (Pani et al. 2012), and the formation of coelomic cavities from an endomesoderm region (Green et al. 2013; Kaul-Strehlow and Stach 2013). The patterning mechanism of DV-polarity is also similar, but chordates appear to invert the DV axis (Lowe et al. 2006; Su et al. 2019). Other functionally analogous structures with uncertain homology include a collar nerve cord that follows a morphogenesis process similar to chordate neurulation (Nomaksteinsky et al. 2009; Miyamoto and Wada 2013) but employing divergent genes for neuronal patterning (Kaul-Strehlow et al. 2017). These homologies could even potentially extend to the cis-regulatory machinery (Yao et al. 2016). Despite remarkable efforts in elucidating the genomic complexity and patterning mechanisms in hemichordates, a comprehensive understanding of the hemichordate developmental programmes and how the genome is temporally regulated is still lacking. Because of their life cycle bearing resemblances to both of their closest living relatives, indirect-developing hemichordates provide an opportunity to investigate the impact of genome regulation on the upbringing of echinoderm-like larval and chordate-like bilateral development. This includes assessing whether a comparable developmental period can be recognized in deuterostomes, and to what extent the developmental gene regulatory networks (GRNs) are conserved among different deuterostome groups. While there have been efforts to begin the functional characterisation of cis-regulatory elements in hemichordates such as *Saccoglossus kowalevskii* (Minor et al. 2019; Yao et al. 2016), the current lack of cis-regulatory information for indirect-developing hemichordates, as well as cis-regulatory data spanning the whole genome, precludes further comparisons to assess the degree of conservation between deuterostomes.

To address these questions, we characterised developmental transcriptomes and chromatin accessibility of *Ptychodera flava*, an indirect-developing hemichordate. We discovered that it exhibits a biphasic developmental program, and identified and compared clusters of stage-specific modules of coexpressed genes controlled at the cis-regulatory level by likewise stage-specific classes of transcription factors (TFs). Network analyses revealed highly interconnected TFs orchestrating gene expression and the recruiting of during gastrula. Comparative developmental transcriptomes of hemichordates, echinoderms, and amphioxus showed conserved gene and TF expression during gastrulation that extends to the neurula stages of amphioxus, along with remarkable similarity in larval transcriptomes across the three species. Additionally, network analyses suggest *P. flava* possesses conserved interactions of transcription factors necessary for the development of echinoderm endomesoderm and chordate axial mesoderm, including conserved cis-regulatory elements of the FoxA TF that is central to the two GRNs. Integration of gene expression data with ancestral linkage group (ALG) data revealed that gene expression recapitulates the independent evolutionary history of each ALG in each deuterostome lineage. Together with gene age analyses showing a conserved gastrulation and younger genes deployed during larval genes, this suggests a potential role of genome rearrangement during the evolution of larval strategies in hemichordates and deuterostome body plans.

## Results

### Two major transcriptional programmes dominated by different TF classes

Indirect-developing hemichordates go through several morphologically distinct developmental stages. After two days of embryogenesis, the late gastrula of *P. flava* hatch and undergo a series of planktonic, tornaria larval stages that stay in the water column for months before metamorphosing into a worm-like body plan (Fig. 1A). We measured gene expression at sixteen stages, covering the entire *P. flava* life cycle, to understand the dynamics of gene usage during its development (Fig. 1A). Principal Component Analysis (PCA) of these transcriptomes reflects the chronological order of the developmental stages from which they were obtained (Fig. 1B, Supplementary Fig. S1), validating the quality of this data series. Overall, we identified 28413 genes expressed across all stages (Sup. Data 1). Similarities at the whole-transcriptome level suggest two major transcriptional programmes of gene expression: early (embryonic) versus late (larval) that run independently (Fig. 1C). Differential gene expression analysis throughout development identified 23726 dynamically expressed genes (hereafter referred to as stage-specific genes), which represent 83% of all genes detected and can be grouped into twenty-two clusters based on their temporal gene expression profiles (Fig. 1D, Sup. Data 1). Several of these gene clusters clearly correspond to their deployment at different developmental phases (e.g. clusters 1-2 match to maternal factors used in early embryogenesis; clusters 6-8 are activated during embryogenesis; clusters 15-19 are generally deployed during the larval phase; clusters 20-22 are predominantly utilised during metamorphosis). We noted that genes expressed at the Muller stage (the newly hatched tornaria) include clusters deployed during embryonic (clusters 12-13) and later larval stages (clusters 15-17 and 19). This observation suggests that Muller stage represents a turning point between embryonic and larval transcriptional states, since its transcriptomic profile is essentially employing embryonic-specific genes, but starting to shift to a ‘larval-like’ transcriptomic programme. Gene Ontology (GO) enrichment analysis of these clusters revealed major developmental processes taking place at the time of peak expression (Table 1). For example, we found GO terms associated with cell cycle divisions and activation of transcription during early divisions, morphogenetic pathways during gastrulation, and formation of body/larval-specific structures at late development clusters (Sup. Fig. S2, Sup. Data 2). These results corroborate our understanding on the temporal controls of transcriptional programmes orchestrating different developmental processes.

**Figure 1:**
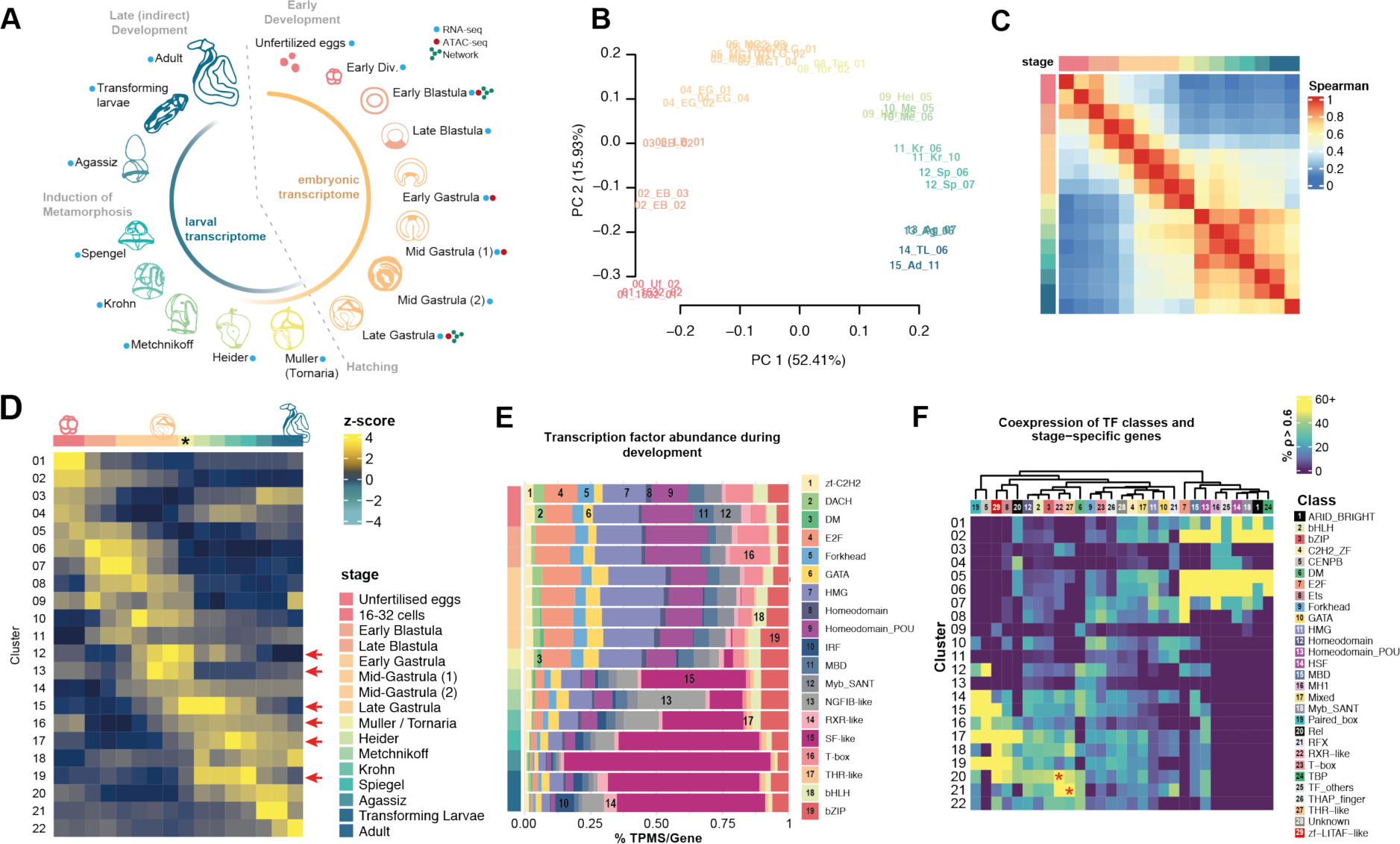
Transcriptomics of *P. flava* during development. A: Life cycle of *P. flava* and sample collection. Grey dashed line represents the time of hatching, and inner coloured lines represent the duration of the embryonic and larval transcriptome. B: PCA analysis of the developmental transcriptomes. C: Two major transcriptional programmes (phases) during development: embryonic and larval development. Pairwise Spearman correlation of transcriptomes of each stage (quantile-normalised, averaged replicates). D: 22 clusters of dynamically expressed genes among the 16 stages. Hierarchical clustering of stage-specific genes by peak expression throughout development (for visualisation, a random subset of 200 genes). E-F: The two phases are orchestrated by distinct sets of TF classes. E: Relative expression (transcripts per million per gene) of each major expressed class during development. F: Coexpression of TFs and stage-specific genes. For a given TF class X and cluster Y, fraction of X genes with a spearman correlation above 0.6 with the average z-score profile of genes in cluster Y.

**Table 1:** Description of differentially expressed, stage-specific gene clusters.

| Cluster ID | Cluster size | Time of expression | Description based on GO terms |
| --- | --- | --- | --- |
| 1 | 1671 | Early Divisions | Protein (de)ubiquitination & phosphorylation, cell response to starvation, DNA methylation |
| 2 | 1352 | Early Divisions | RNA processing, histone modification, DNA repair, sister chromatid cohesion, nuclear mRNA surveillance, initiation of transcription |
| 3 | 879 | Early Div., Agassiz, Adult | Phospholipid catabolic process, regulation of developmental growth, transport & catabolism of biomolecules |
| 4 | 696 | Early Divisions, Larval stages | Lipid homeostasis, vesicle and endosome organisation, sterol biosynthesis |
| 5 | 508 | Early Div., Early Blastula | Positive regulation of Wnt pathway, protein polyubiquitination, spindle assembly, membrane fusion, mitochondrial fission, regulation of gene expression, histone acetylation |
| 6 | 1436 | From Early Div. to Mid Gastrula | mRNA splicing, RNA processing, DNA replication, centromere assembly, mitosis |
| 7 | 1204 | From Early Blast. to Mid Gastr. | RNA processing, gene expression, histone acetylation, patterning formation |
| 8 | 1247 | From Late Blast. To Mid Gastr. | Telomere maintenance, protein folding, cilium assembly, mRNA splicing, IMP biosynthesis. Vesicle fusion, histone exchange |
| 9 | 226 | Early Div, Mid Gastr., Adult | Response to nitrogen compound, DNA integration |
| 10 | 1038 | Mid Gastrula to Larval stages | glycolipid, IMP, ATP biosynthesis, protein glycosylation, organ morphogenesis & development, cellular respiration |
| 11 | 1296 | Early Blast. To Larval stages | Protein degradation, DNA repair, Wnt pathway, vesicle formation, patterning, respiration |
| 12 | 582 | Late Gastrula to Metschnikoff | Muscle contraction, calcium transport, lipid metabolism |
| 13 | 321 | Late Gastrula, Tornaria | Sphingolipid biosynthesis, cilium assembly, cell projection, sodium transport. Potentially related to axon formation and/or larval cilia |
| 14 | 1539 | Mid Gastrula to Adult | Ribosomal assembly, vesicle transport, exocytosis, actin filament processes. |
| 15 | 943 | Late Gastrula to Agassiz | Axoneme assembly, mesenchyme development, cilium formation. Potentially related to axon formation and/or larval cilia |
| 16 | 1276 | Early Div., Tornaria to Adult | Cilia motility, axon formation and guidance, Smoothened pathway, protein transport |
| 17 | 1512 | Heider to Agassiz, Adult | Signal transduction, ERK cascade, ion transport, actin nucleation |
| 18 | 895 | Heider to Adult | Muscle development, Notch and TLR pathways, thyroid hormone metabolism, proteolysis, DNA integration, apoptosis |
| 19 | 1440 | Heider to Spengel | Ion and phospholipid transport, telomere organisation, glycoside metabolism, netrin pathway |
| 20 | 1379 | Krohn to Adult | Ion transport, signal transduction, nervous system process, cell communication, gluconeogenesis |
| 21 | 1242 | Agassiz to Adult | Transmembrane transport, trans-synaptic signalling, regulation of growth, protein glycosylation, nucleotide biosynthesis, catecholamine metabolism, thyroid hormone metabolism |
| 22 | 1044 | Transforming larvae, Adult | Apoptosis, regeneration, immune system development, SMAD transduction, steroid biosynthesis, hematopoiesis or lymphoid organ development, TLR pathway |

To understand how the transcriptional programmes are controlled, we focused on the usage of transcription factors (TFs) during *P. flava* development. For an approach to identify the underlying gene expression regulatory machinery, we classified the transcription factors into major classes (following animalTFDB and automated orthology relationships with vertebrates) and looked at their expression throughout development (Sup. Data 2, Sup. Data 3). We observed two distinct programmes of TF composition, with a high prevalence of T-box, Pou and High Mobility Group (HMG, which include Sox) TFs that are maternally deposited and used at early stages, and SF-like, Myb-SANT and bZIP TFs taking over at larval stages (Fig. 1E), and with a higher number of Homeobox TFs expressed at later stages (Supplementary Fig S3). The timing of these TF classes generally correspond to their roles in development in other species. For example, the Pou and T-box (including Brachyury) classes, mediate pluripotency and cell fate specification in early stages and gastrulation (Nichols et al. 1998; Niwa, Miyazaki, and Smith 2000; Babaie et al. 2007; Osorno and Chambers 2011), while SF-like classes related to steroidogenic signalling and gonad maturation function at later stages (Mangelsdorf et al. 1992). Additionally, several TF classes exhibit high agreement in coexpression with stage-specific gene clusters throughout development, supporting that the two distinct developmental programmes of embryonic and larval development are controlled by different TF classes (Fig.1F). Notably, RXR (class 22) and Thyroid hormone (THR) (class 27) pathways are involved in metamorphosis in a variety of animals (Xuedong Wang, Matsuda, and Shi 2008; Shao et al. 2017), and their co-expression with clusters 20-21 (red asterisks in Fig. 1F), genes predominantly expressed during metamorphosis, suggests the similar role in *P. flava*. These results indicate that the two major developmental programmes of *P. flava* are controlled by two sets of TFs to perform stage-specific processes known in other deuterostomes.

### Gastrulation retains the oldest transcriptome in *P. flava*

To further delineate the evolutionary history of the different transcriptional programmes, we sought to investigate the evolutionary conservation of the gene expression programme underlying *P. flava* development. To this end, we firstly characterised the gene age of the whole annotated genome of *P. flava*. We assigned groups of orthology (orthogroups) using OMA (Altenhoff et al. 2021) and a comprehensive set of metazoan species, with an emphasis on deuterostomes and ambulacraria (Fig. 2A, Sup. Data 4). In turn, we assigned a gene age to every orthogroup by Dollo parsimony (Csurös 2010), resulting in ancient genes present in all species and younger genes found only in *P. flava* (see Methods) (Fig. 2A).

**Figure 2:**
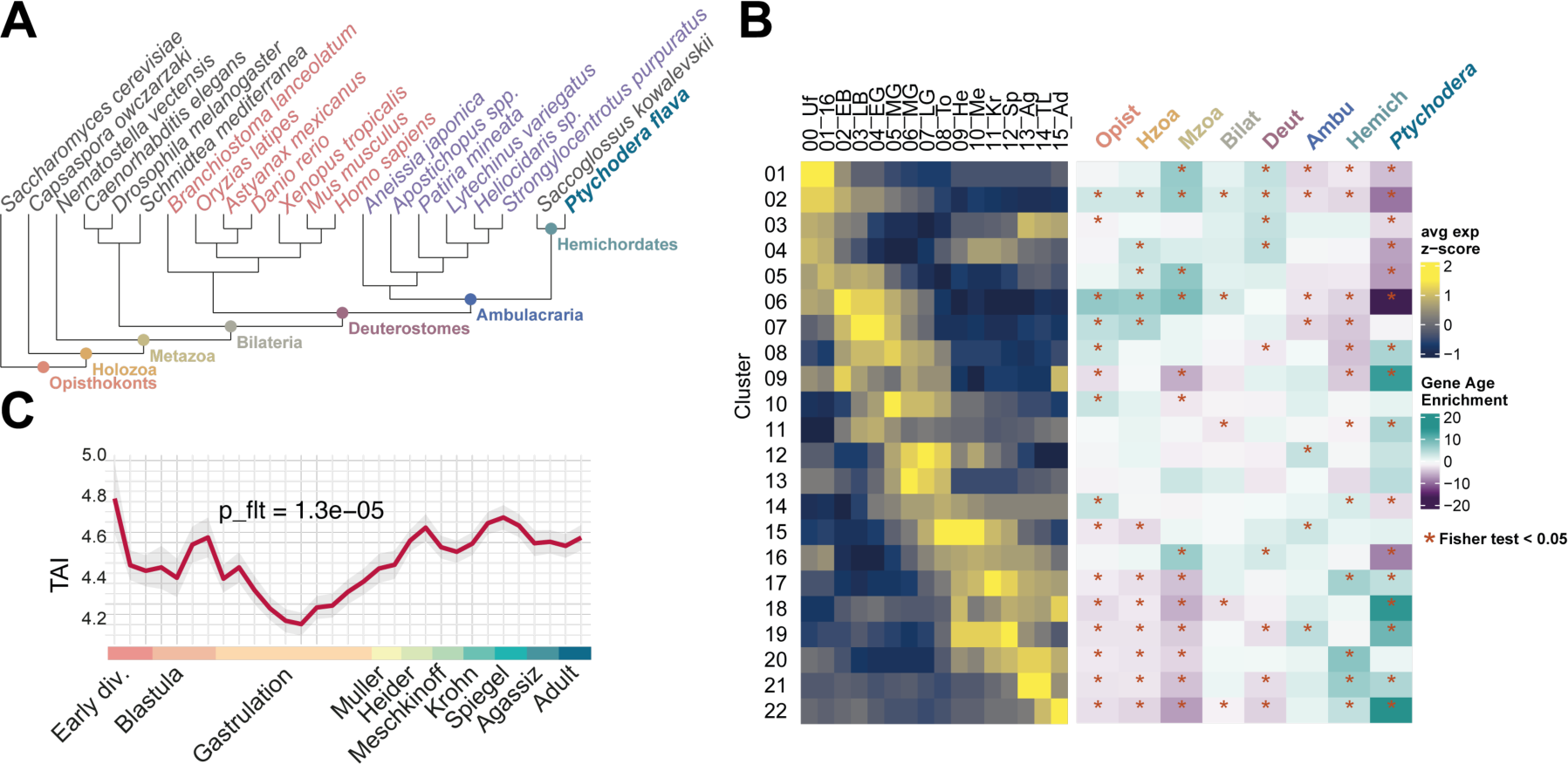
Gene Age of the developmental transcriptional programme in *P. flava*. A: Tree of species used for the orthology and gene age assignment analysis. B: Age of the 22 clusters: (Left) Schematics of average gene expression for each stage-specific cluster (summary of Fig. 1D); and (Right) gene age enrichment of stage-specific clusters. C: Transcriptomic Age Index (TAI) throughout the development of *P. flava*.

We first calculated gene age enrichment for every cluster of stage-specific genes, by comparison with gene age of all annotated *P. flava* genes. We found that clusters of early stage genes were slightly enriched in genes from Opisthokonta all the way up to Bilateria, but slightly depleted in ambulacraria- or *Ptychodera*-specific genes, and an opposite trend was found in clusters of later stages with minor enrichments in ambulacraria-, hemichordate- or *Ptychodera*-specific genes (Fig. 2B). This result suggests that older genes tend to be expressed during embryogenesis, while younger genes are more probably deployed for larval development and the metamorphosis process leading to the adult body plan. This matches reports of similar trends in echinoderms (Li et al. 2020). To further evaluate the transcriptome age among different developmental stages, we used the same approach to calculate the Transcriptomic Age Index (TAI) (Domazet-Lošo and Tautz 2010), taking into account both gene age and expression level of all the genes of *Ptychodera* at a given developmental stage. TAI analysis identified the mid-to-late gastrula stage as the oldest transcriptome relative to other stages (Fig. 2C). Overall, this suggests that embryogenesis tends to utilise older genes with gastrulation being a more conserved stage of development.

### Cis-regulation and gene regulatory networks during hemichordate gastrulation

For a better understanding of how the gene expression programme is controlled at the cis-regulatory level before and during the conserved gastrulation period, we identified open chromatin regions in the genome using Assay for Transposase-Accessible Chromatin sequencing (ATAC-seq) (Buenrostro et al. 2013) at four time points: early blastula, early-, mid-, and late gastrula (Fig. 1A, indicated with the red dots). We identified 82807 open chromatin regions (OCRs) that are potential cis-regulatory elements (REs) throughout gastrulation in *P. flava*, mostly located in the ±20kb vicinity of genes, specifically within intergenic regions or in introns, followed by within promoters (Sup. Fig. S4 A,B, Sup. Data 5,6). Binding motifs of trans regulators such as developmental TFs appear more frequently outside promoters and in distal elements (Figure S5 Supp). Nearly eighty percent of the annotated genes have at least one OCR associated, with more than fifty percent associated with two or more OCRs (Fig. S4C Supp). We compared the signal intensity of the ATACseq data at OCRs and RNAseq levels at their associated genes and observed a trend of OCR openness in accordance with the expression level of nearby genes (Sup. Fig. S4D), suggesting cis-regulation generally occurs as transcriptional activation. Genes annotated as trans-developmental (with GO terms in transcription or development, as similarly defined in (Marlétaz et al. 2018), see Methods) tend to be associated to cis-regulatory elements, with more OCRs per gene than housekeeping genes (Sup. Fig. S4 E,F,G). This is in agreement with previously observed trends towards complexification of cis-regulatory architecture in genes orchestrating metazoan development (Irimia et al. 2012; Marlétaz et al. 2018; Bogdanović et al. 2012).

To identify temporal patterns at the chromatin level, we performed differential accessibility analysis and identified 15946 dynamically accessible OCRs throughout gastrulation, which were categorised into six clusters (Fig. 3A). These six clusters were enriched with specific DNA binding motifs of major TF classes that correspond to their roles in controlling gene expression during development in other metazoan lineages (Fig. 3B, Figure S6 Supp). Motifs of blastula-specific OCRs (01_EB) show little overlap with gastrula-specific OCRs (03_EG_MLG to 06_LG) (Fig. 3B). For example, motifs of T-box transcription factors (including Brachyury, Eomes, Tbr and Tbx) are enriched in the OCRs at the early blastula and early gastrula stages. On the other hand, Sox (HMG) binding motifs are enriched in OCRs after gastrulation occurs. The dynamic of motifs enriched in stage-specific OCRs is consistent with the transcriptome data, in which T-box and HMG (including Sox) factors are highly expressed during embryogenesis, with T-box predominantly present before gastrulation and HMG more abundant during gastrulation (Fig. 1E). Other examples of stage-specific OCRs include binding motifs of E2F and the p53/63/73 family that are related to cell division, and of Snail, Dlx, Fox, Gsc and Pax that are known to express in different germ layers during *P. flava* gastrulation (Fig. S6 Supp) (Su et al. 2019; Harada et al. 2001; E. Röttinger and Martindale 2011; Taguchi et al. 2000). We crossed these data with early stage-specific genes and also found binding motifs for early transcription factors, like T-box, E2F and HMG (Fig. 3C), correlating with the early factors found based on direct ATACseq data clustering (Fig. 3B). This concordance between the binding sites of stage-specific OCRs and the binding sites in the proximity of stage-specific genes, suggests that potential regulatory features of stage-specific genes can be inferred based on their surrounding cis-regulatory landscape.

**Figure 3:**
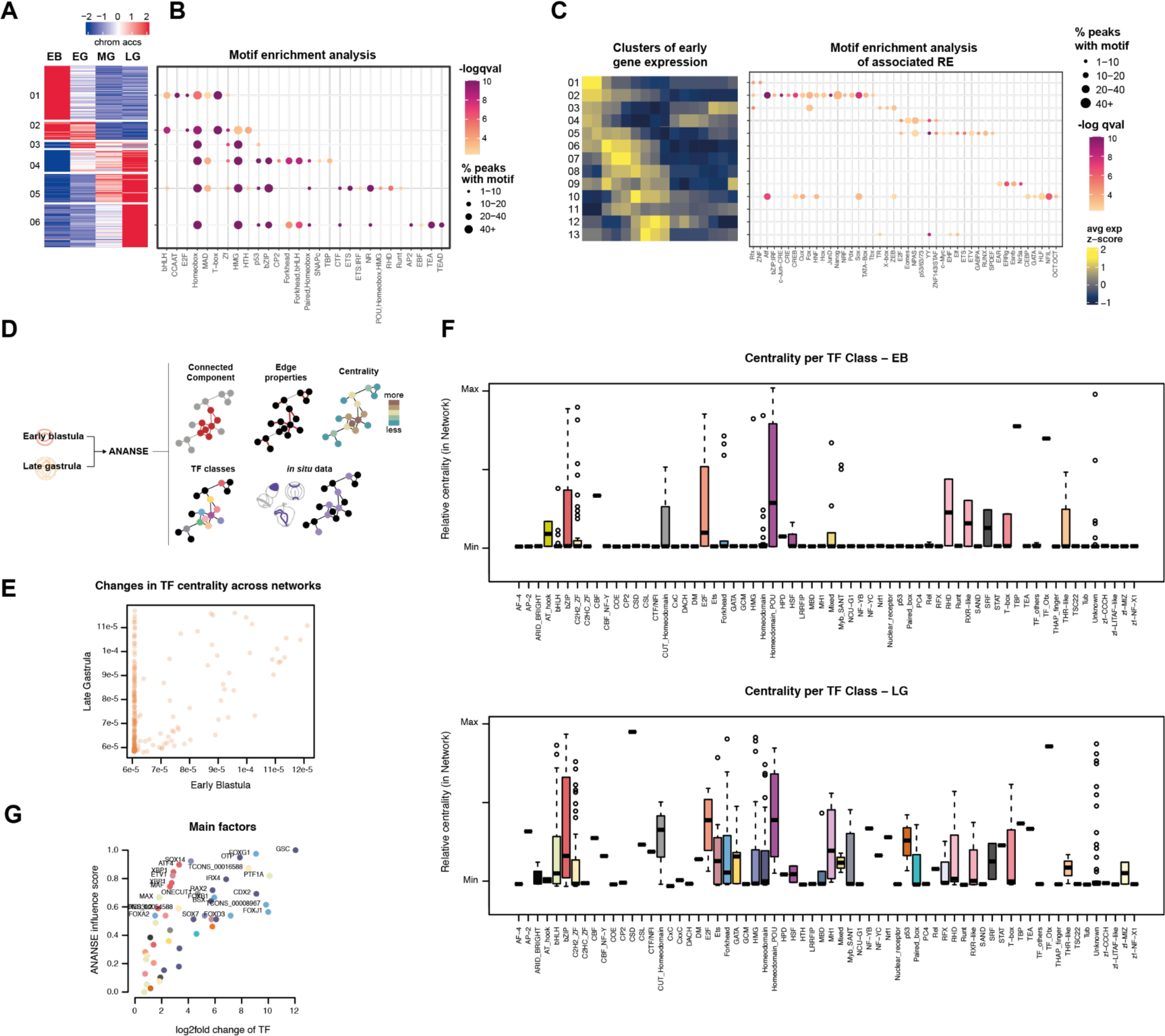
Cis-regulation of *P. flava* during gastrulation. A: Clustering of dynamically accessible OCRs over time. B: Motif enrichment analysis of each cluster of OCRs in panel A. Dot plot for HOMER logqvalue (colour) and fraction of peaks with motif (dot size). C: Motif enrichment analysis of OCRs associated with early stage-specific genes: (Left) Schematics of average gene expression for each stage-specific cluster (summary of Fig. 1D); (Right) Dot plot for HOMER - log(p.adj) (colour) and fraction of peaks with motif (dot size). D: Schematic of the ANANSE analysis for networks of cis-regulatory interactions. E: scatter plot of TF centrality in each of the networks. X axis : centrality in Early Blastula network ; Y axis: centrality in Late Gastrula network. F: Distribution of centrality in network for all the TF classes in the Early Blastula network (up) and the Late Gastrula network (down). G: Scatter plot of main factors driving the transition as calculated by ANANSE influence. X axis: logfold change of gene expression. Y axis: ANANSE influence score. Top 20 TFs are labelled with their eggNOG predicted gene name.

Processes in development do not result from independent actions of individual genes but are rather driven by complex interactions between many genes constituting developmental gene regulatory networks (GRNs). To assess putative GRNs in *P. flava*, we used ANANSE (Xu et al. 2021), a computational tool that uses cis-regulatory and gene expression data to generate graphs and predict TF-target gene interactions. These graphs are composed of nodes (the TFs and target genes) and edges (the interactions between TFs and target genes, whose score values are calculated using gene expression, OCR accessibility, distance between OCR and its corresponding transcription start site, and motif enrichments within the OCR). We generated two independent graphs, one of early blastula (EB) and one of late gastrula (LG) (Fig. 1A, 3D) (Sup. Data 7,8). The number of genes represented in both graphs is similar, around 8000 genes (Sup. Fig. S7 A), with many more TF genes in LG (242 TFs) than in EB (79 TFs) (Sup. Fig. S7 B).

Consistently, we observed a net gain of connectivity in LG compared to EB, as seen in differences in TF centrality (Borgatti 2005) (how much a given gene is embedded within each GRN, related to the number of connections to other genes) (Fig. 3E) and other network metrics (Supp Fig. S7C-G). This net gain of connectivity is attributed to a higher TF class diversity in late gastrula (Fig. 3F). Overall, TF classes showing higher centrality correspond to the dominantly expressed TFs at the respective stage (Fig. 1E,F). For example, E2F and Pou classes show high centrality in the EB network, and bZIP, ETS, GATA, Homeobox and Forkhead factors show higher centrality in the LG network. This suggests a different set of core TFs controlling the GRNs at each stage. TFs common between the two networks are enriched in terms of regulation of gene expression, which we found reassuring of our analysis, while stage-exclusive TFs of LG are enriched in developmental processes (Sup. Fig. S8 A). The genes that are being controlled by these networks are also different. They share 499 target genes (Sup. Fig. S8 B), with overall overlapping GO terms in the two stages but higher enrichments for cell migration and organogenesis in LG-specific target genes (Sup. Fig. S8 C); when looking at functional categories, we see an enrichment in genes annotated with signal transduction in gastrulation (Sup. Fig. S8 D). Interestingly, more than 50% of genes from clusters specific to larval stages were detected inside the LG graph compared to EB (Sup. Fig. S8 E). These observations suggest a different core of TFs controls genes related to organogenesis and signalling during gastrulation in *Ptychodera*, including genes that are more highly expressed after gastrulation.

To further inspect whether the GRNs can be subdivided into circuits that mediate germ layer formation, we investigated the position of *P. flava* genes that are known to show germ layer-specific expression at the LG stage in the LG network (Sup. Data 9, Sup. Fig. S9 A,B,C). Within those circuits, we noticed that TF genes showing high centrality tend to express broadly, while low centrality genes often display restricted expression patterns in the respective germ layer. For example, xbp1 (Eric Röttinger et al. 2015), *soxb2* and *otx* are the most connected genes within the ectoderm circuit (Sup. Fig. S9 D, E) and are known to express in broad ectodermal domains at the LG stage (Sup. Fig. S9 E). Similarly, genes show high centrality in the mesoderm network (e.g., *foxg* and *six3*) and endoderm circuit (e.g., *soxb1a* and *foxa*) (Sup. Fig. S9 D, E) are expressed broadly in the mesoderm-derived protocoel and endoderm-derived archenteron of the LG embryos, respectively (Sup. Fig. S9 E). Genes that are known to express in restricted regions in the corresponding germ player, such as *xlox* (posterior ectoderm; (Ikuta et al. 2013)), *foxq2* (anterior ectoderm; (E. Röttinger and Martindale 2011)), *msx* (dorsal mesoderm; (Eric Röttinger et al. 2015)), and *tsg* (anterior endoderm; (Eric Röttinger et al. 2015)) show low centrality in the respective circuit (Sup. Fig. S9 D). Other examples of TF genes with low centrality displaying restricted germ layer-specific expression patterns are shown in Sup. Fig. S9 E. The agreement between spatial expression patterns of TF genes and their hierarchical positions in GRNs coincides with the expected roles of TFs in specifying germ layers and the subdomains constituting the specific germ layer. We also observed that peripheral TFs tended to be more associated with organogenesis and other tissue-restricted processes, compared to more central TFs (Sup. Fig. S9 F). We finally used ANANSE influence to detect the factors responsible for the transition between the network states of Early Blastula and Late Gastrula (Xu et al. 2021), and found a number of well-known developmental TFs with known roles in germ layer-specific fate induction such as Cdx, Irx4, FoxG2 and FoxA (Fig. 3G, Sup. Fig. S9 G).

Altogether these results suggest a more interconnected GRN at late gastrula by addition of factors and targets to the blastula GRN, including genes to be deployed later in development during larval stages. The highly connected GRN in late gastrula is achieved by employing diverse TF classes that exhibit higher self-regulation. This feedback property is expected to stabilise distinct regulatory states for germ layer specification. Additionally, TF genes exhibiting high centrality correspond to observations at their spatial expression patterns, consistent with their hierarchical roles within the individual germ layer networks.

### Comparative transcriptomics between *P. flava* and other deuterostomes

To globally assess the degree of conservation in developmental programs among the three major deuterostome groups, we performed pairwise comparisons of developmental transcriptomes between *P. flava* and two other invertebrate deuterostome species: the purple sea urchin *Strongylocentrotus purpuratus* and the lancelet (amphioxus) *Branchiostoma lanceolatum*. Whole transcriptome, pairwise comparisons between *P. flava* and sea urchin (Tu et al. 2012; Tu, Cameron, and Davidson 2014) using 5604 one-to-one orthologues revealed high similarities at the gastrula stages (Fig. 4A, left panel; Sup. Fig. S10A). We noted that the hemichordate early gastrula transcriptome bears high resemblance to that of the sea urchin mesenchyme blastula, which represents the initial gastrulation stage when the primary mesenchyme cells ingress into the blastocoel of sea urchin embryos (Hardin and Cheng 1986). We found genes driving similarities between these stages were enriched in terms related to regulation of gastrulation, with NODAL as one example (Sup. Fig. S10B). We also observed high similarities at larval stages of the two ambulacrarian species, both at the pairwise-comparisons of developmental stages and at the number of overlapped gene families (Sup. Fig. S10C). In the latter comparison, we observed gene families common between pairs of clusters of post-gastrulation genes were enriched in functions of signal transduction and secondary metabolism (Fig. 4C), which matches with the described functions for post-gastrulation clusters in Ptychodera (Fig.1D, Table 1). Interestingly, we found similar trends when comparing *P. flava* to *Anneissia japonica*, an echinoderm from the class Crinoidea that is sister-group of the rest of echinoderms (Li et al. 2020) (Sup. Fig. S11). The high similarities between ambulacrarian larval transcriptomes supports the hypothesis that hemichordate and echinoderm larvae are homologous and the dipleura-type larva is an ancestral trait of ambulacraria (Tagawa 2016). These results point to a similar direction as Fig. 2B, supporting the idea that the last common ancestor of ambulacraria (LCA) possessed larval development.

**Figure 4:**
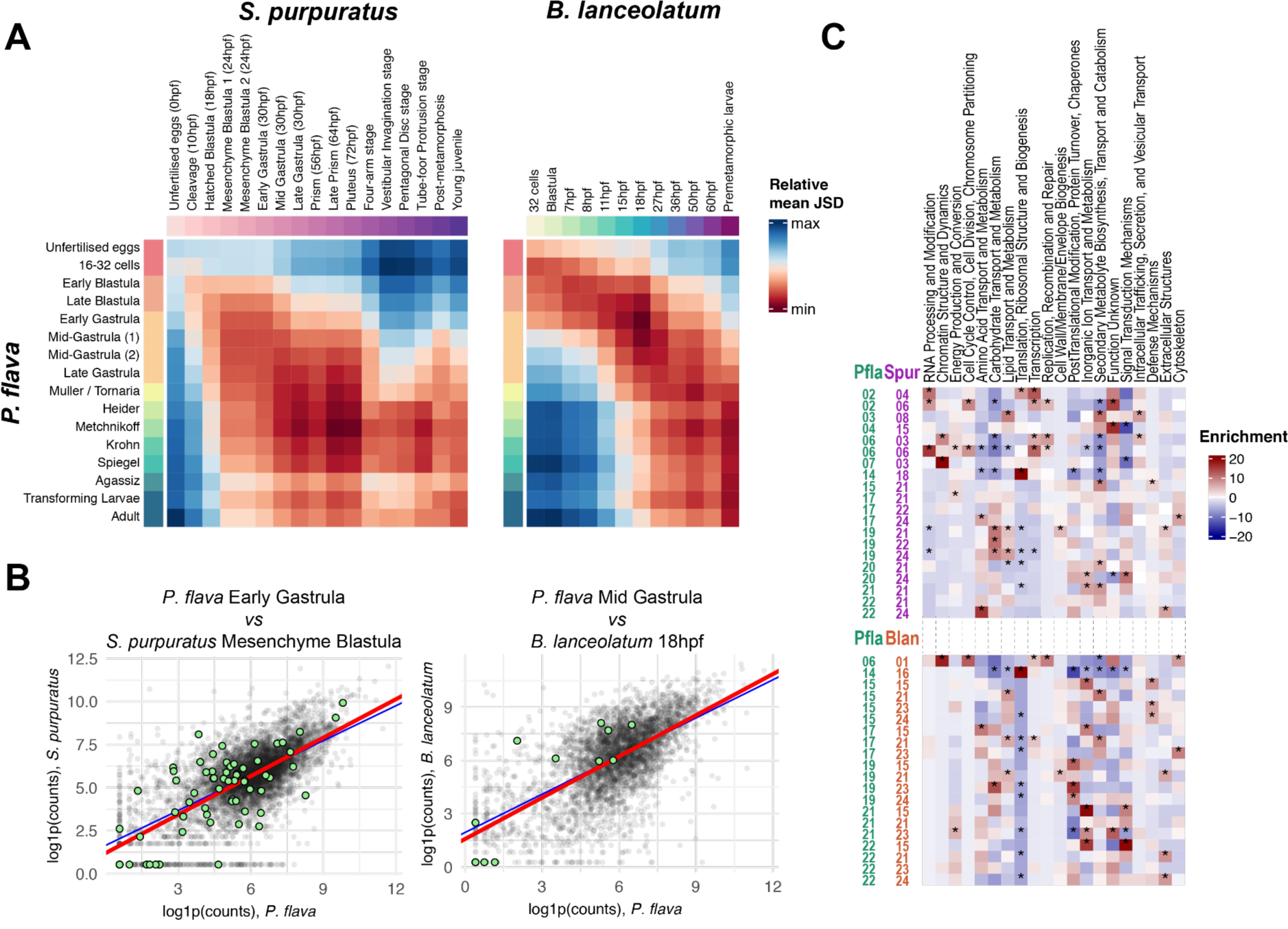
Comparative transcriptomics of Ptychodera and other deuterostomes. A: (Left) mean Jensen Shannon Distance (bootstrap 1000 iterations) of pairwise-comparisons between P. flava and sea urchin developmental transcriptomes. (Right) Same analysis as A for *P. flava* and the amphioxus developmental transcriptomics. B: Scatter plot of the expression (log1p counts) of 1-to-1 orthologs between *P. flava* early gastrula and *S. purpuratus* mesenchyme blastula (left) and between *P. flava* mid-gastrula and amphioxus at 18h (right). Green highlights the genes found expressed in these pairs of stages and not others; i.e. the “unique genes” responsible for the similarities observed (see Methods). C: Functional Category Enrichment Analysis between pairs of overlapping developmental clusters (see Sup. Fig.; Methods) of *P. flava*, *S. purpuratus*, and *B. lanceolatum*. Asterisks indicate Fisher Test p < 0.05.

The pairwise comparison between *P. flava* and amphioxus (Marlétaz et al. 2018) also showed resemblance of the gastrula transcriptomes (Fig. 4A, right panel; Sup. Fig. S12A). Intriguingly, this similarity extends to the amphioxus neurula stages. As to the genes driving similarities at these stages, we found genes responsible for regulation of neuron migration, cell fate specification and commitment, and epithelial-to-mesenchymal transition, including homologs of Wnt, Goosecoid, Oligodendrocyte transcription factor OLIG3, and HELT transcription factor, involved in neuronal specification (Nakatani et al. 2007; Miyoshi et al. 2004) (Fig. 4B, right panel; Sup. Fig. S12B,C). We observed a very similar trend when comparing the transcriptomes of *P. flava* with the developmental transcriptomes of another amphioxus species, *Branchiostoma floridae* (Haiyang Hu et al. 2017) (Sup. Fig. S13). The observation that amphioxus neurula stages are transcriptomically similar to hemichordate gastrula likely reflects the exact timings of comparable developmental events; for example, neurogenesis, as well as separation of mesoderm and endoderm occurs at the mid-gastrula in *P. flava* and during neurulation in amphioxus (Lin et al. 2016; Carvalho et al. 2021). As the transcriptomes of amphioxus neurula stages also show the highest similarity when compared to those of several vertebrates (Marlétaz et al. 2018), this period (gastrula stages of ambulacrarians and gastrula+neurula stages of chordates) may represent the putative phylotypic stage of deuterostomes. An additional unexpected observation is the transcriptomic similarity between hemichordate and amphioxus larval stages (both *B. lanceolatum* and *B. floridae*), when these species start to feed (Fig. 4A, right panel; Sup. Fig. S12 A, Sup. Fig. S13). We also observed these similarities during late development when looking at common gene family usage (Sup. Fig. S12C) in pairs of stage-specific clusters and their enriched functional categories (Fig. 4C). These were genes enriched in functional categories such as primary and secondary metabolism, and signal transduction (Fig. 4C), which align with GO terms found in pairwise comparisons of larval-specific genes such as ‘gland/organ formation’ (Sup. Fig. S12 D). We observed similar trends in the pairwise comparisons and the orthogroup overlap estimations when using other orthology methods such as OrthoFinder (Emms and Kelly 2015) (Sup. Fig. S14A-F) or BLAST reciprocal best hits (Altschul et al. 1997) (Sup. Fig. S14G-H). Overall, these results suggest that despite the morphological disparity in their larval forms, similar transcriptional programs are utilised by hemichordate and amphioxus larvae.

### Conservation of cis-regulatory mechanisms between deuterostomes

To better understand the underlying mechanisms driving these similarities at the gene expression level, we narrowed our comparison to the transcription factor programme of the three species. We looked at the TF composition of the developmental stages of these three species and noticed similar uses of Pou, HMG and T-box before larval development, and bZIP, Homeobox and SF-like during and after larval development (Sup. Fig. S15). Consistently, these trends roughly match similar observations from published data in ecdysozoa (Adryan and Teichmann 2010), with the early and late deployment of HMG and bZIP factors respectively. We subset our pairwise transcriptomic comparisons to common one-to-one orthologous TFs and retrieved similar trends (Sup. Fig. S16A). In these comparisons we found one TF gene (FoxA) as a contributor to similarities between *P. flava* and amphioxus gastrulation as well as highly expressed during similar stages of development of *P. flava* and sea urchin (Fig. 5A). FoxA is one of the TFs showing highest centrality in the *P. flava* gastrula network (Sup. Fig. S16 B) and the most central of the Forkhead family (Sup. Fig. S16 C), and is found among the TFs regulating the transition between the EB and LG cis-regulatory landscape according to ANANSE influence (Sup. Fig. S16 D). Given its relevance in the predicted *Ptychodera* networks and its similar temporal expression profile between species (Fig. 5B), we wondered whether we could identify cis-regulatory interactions from deuterostome developmental GRNs involving FoxA orthologues in *Ptychodera*. For this we queried the late gastrula network of *Ptychodera* for orthologues of the endomesoderm kernel GRN and the axial mesoderm kernel that are known to be conserved in echinoderms and chordates, respectively (Hinman et al. 2003; Yu et al. 2007). We found the majority of cis-regulatory interactions of the endomesoderm kernel, as well as the axial mesoderm cis-regulatory network, were also retrieved in *Ptychodera*, with FoxA at the centre of the two (Fig. 5C, Sup. Data 10, Sup. Fig. S9). This hints at the conservation of stable subcircuits that underpin developmental programmes (E. H. Davidson 2011, 2010). This result suggests that the conservation of the endomesoderm kernel could be pushed further to the common ancestor of Ambulacraria, reinforcing the notion that part of this kernel already existed in the bilaterian ancestor (Boyle, Yamaguchi, and Seaver 2014). The identification of the cis-regulatory interactions for specifying chordate axial mesoderm in *P. flava* suggests that the genetic circuit predates the emergence of novel structures in chordates, such as the axial mesoderm-derived notochord. Overall these results suggest the potential to establish cis-regulatory interactions of chordate networks could date back to the common ancestor of deuterostomes.

**Figure 5:**
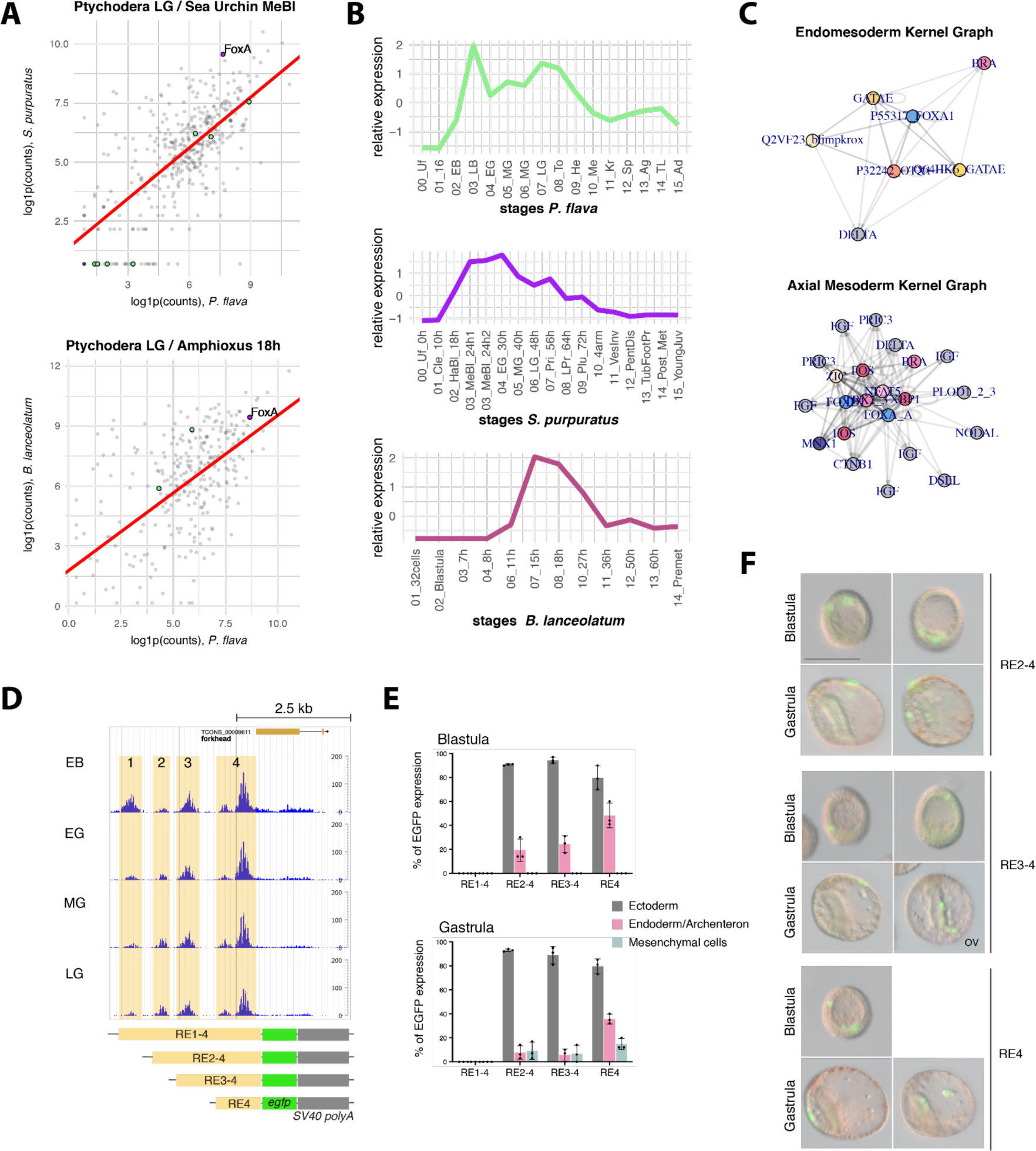
Conservation of the cis-regulatory mechanisms of the TF programme in deuterostomes. A: (Above) Scatter plot of log1p counts of 1-to-1 orthologous transcription factor genes at *Ptychodera* late gastrula and sea urchin mesenchyme blastula stages, highlighting the ones markedly expressed at these stages (green) as well as FoxA as highly expressed (purple). (Below) Similar analysis between *Ptychodera* late gastrula and amphioxus neurula (18h), only FoxA (purple) is also retrieved as markedly expressed in this pair of stages. Green highlights the genes found expressed in these pairs of stages and not others; i.e. the “unique TFs” responsible for the similarities observed (see Methods). B: Temporal expression profiles of FoxA in *Ptychodera*, sea urchin, and amphioxus throughout development. C: Cis-regulatory interactions of the *P. flava* orthologues of the genes constituting the echinoderm endomesoderm kernel (top network) and the chordate axial mesoderm GRN (bottom network). D: Upper panel, ATAC-seq data showing 4 putative REs (shaded in yellow) upstream of the *P. flava* foxa gene at 4 different developmental stages. RE3 and RE4 are accessible at all 4 stages; RE1 accessibility decreases, while RE2 accessibility increases during gastrulation. Lower panel, illustrations of the four reporter constructs assayed in sea urchin embryos. E: Percentages of EGFP expression domains driven by the four reporters at the blastula and gastrula stages. F: EGFP expression driven by the indicated reporter constructs at the blastula and gastrula stages. All embryos are shown in the same scale (scale bar: 100 µm). Unless otherwise indicated, the gastrulae are viewed from the left side with ventral (oral) to the left. OV: oral view.

To assess the degree of conservation of the cis-regulatory information of FoxA, we generated reporter constructs covering one to four potential REs upstream of the *Ptychodera foxA* gene (Fig. 5D), the expression of which is initiated at the early blastula stage (Fig. 5B) in the presumptive endomesoderm (Su et al. 2019; Taguchi et al. 2000). Using sea urchin embryos as a surrogate system, we showed that these REs are able to drive specific expression patterns, with RE4 as a tissue-specific activator and RE1 as a general repressor (Fig. 5E-F). The repressive role of RE1 is consistent with the decrease of its openness after the early blastula stage (Fig. 5D), when the gene starts to express zygotically (Fig. 5B). These results validate the functional roles of OCRs revealed by ATAC-seq, and suggest similar regulatory inputs to the *foxA* gene during sea urchin and hemichordate gastrulation. Overall these results allowed us to reconstruct potential cis-regulatory interactions of the common ancestor of deuterostomes.

### Gene expression under the context of deuterostome chromosome evolution

The *foxa* gene is part of the pharyngeal gene cluster, a group of genes linked together throughout metazoan evolution (Kapli et al. 2021; Simakov et al. 2015; X. Zhang et al. 2017). Previous studies supported the idea of conserved synteny in the pharyngeal cluster of hemichordates (Simakov et al. 2015), which we corroborated in our companion manuscript studying the genome architecture of *Ptychodera flava* and *Schizocardium californicum* under the context of bilaterian evolution (Lin et al., in prep.). Provided the conserved expression of FoxA during development shown above, we wondered if there was an association between gene expression and gene clustering in the genome. To explore whether pharyngeal genes are co-regulated when clustered, we analysed the temporal expression profiles of pharyngeal orthologous genes using available transcriptomic datasets (Gaiti et al. 2015; Gildor et al. 2019; Haiyang Hu et al. 2017; Leclère et al. 2019; Tu, Cameron, and Davidson 2014; J. Wang et al. 2020; Xiaotong Wang et al. 2014; G. Zhang et al. 2012; L. Davidson, Kerr, and West 2012). In the four deuterostome species we examined (BFL, PFL, SPU and PMI), the overall expression profiles were generally similar, with maternally loaded *xrn2*, *slc25a21a* and *mipol1* transcripts and activation of *nkx2*, *msx* and *foxa* genes immediately before or during gastrulation (Fig. 6A-B). One exception was the amphioxus *msx* gene, the expression of which was not synchronised with *nkx2* genes; coincidently, the location of this gene is 1.8 Mb away from the cluster (Fig. 6A). In the two examined spiralians (CGI and PYE), the *nkx2.1*, *nkx2.2* and *msx* genes were not activated simultaneously, consistent with their lack of close association (Sup. Fig. S17 A-B). Activation of the pharyngeal orthologous genes was less synchronised in jellyfish (CHE) and sponge (AQU), and no syntenic relationships were observed in the current versions of the genomes (Sup. Fig. S17 C-D). These results support the idea that clustering of the pharyngeal genes likely contributes to their co-regulation. Additionally, we noted that the transcript level of *xrn2* shows an opposite trend compared to those of *nkx2*, *msx* and *foxa* genes in deuterostomes. As such, the *xrn2* transcript level decreases while levels of *nkx2*, *msx* and *foxa* mRNAs increase during gastrulation. Xrn2 exoribonuclease is known to promote transcription termination and to degrade several types of RNA, including pre-mRNA (L. Davidson, Kerr, and West 2012; Hori, Yamamoto, and Obika 2015; M. Wang and Pestov 2011). Based on the complementary expression profiles, we hypothesise that *nkx2*, *msx* and *foxa* transcript levels may be down-regulated post-transcriptionally by Xrn2 at early embryonic stages. The synchronisation of the pharyngeal gene expression is thus likely to be controlled at the transcription level due to gene clustering and at the post-transcriptional level due to the 5’ to 3’ exoribonuclease activity of Xrn2. Future functional studies will be required to test this hypothesis.

**Figure 6:**
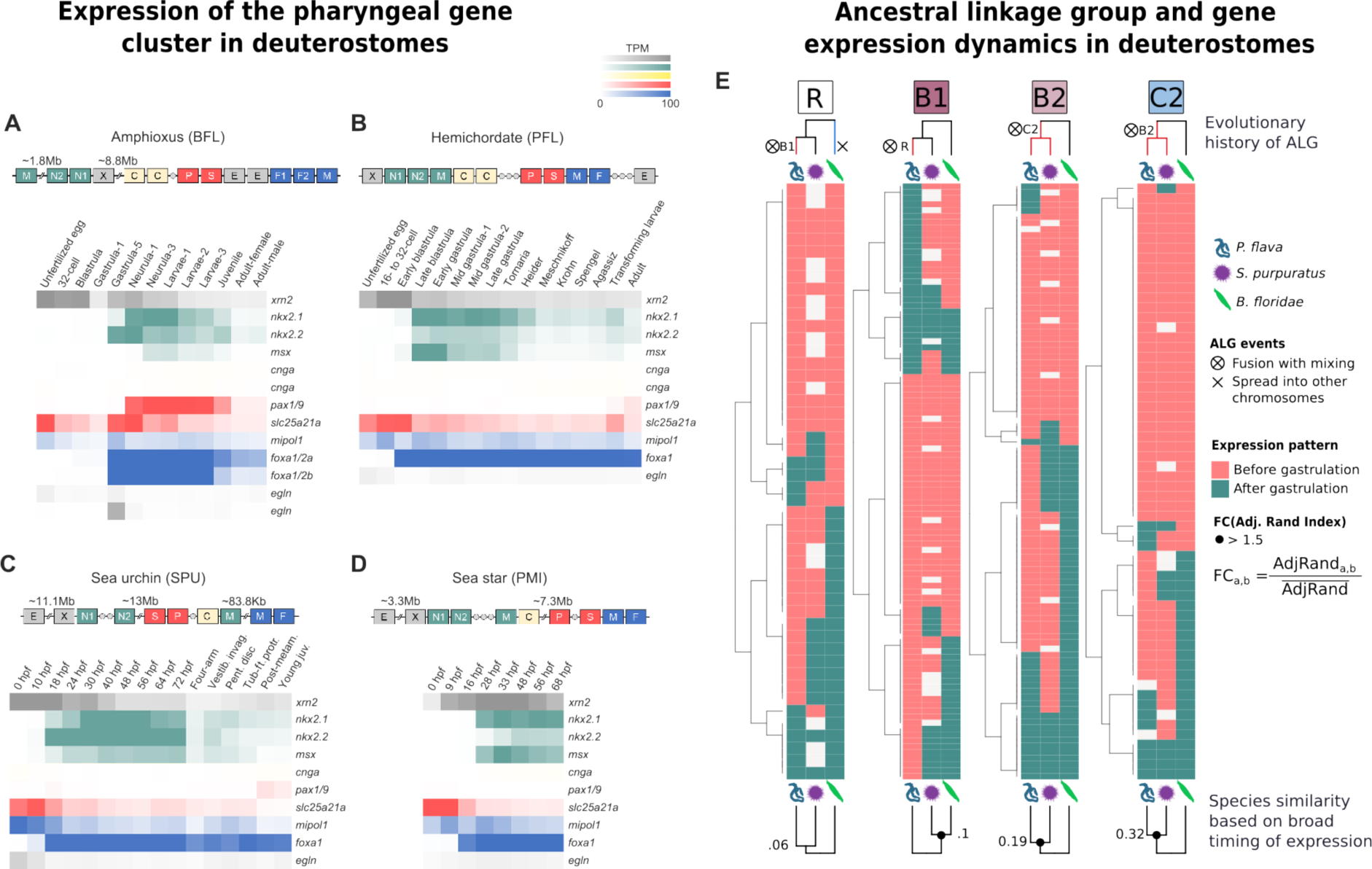
Gene expression and genome architecture during deuterostome evolution. A-D: Expression of the pharyngeal gene cluster genes across four deuterostome species. E: Gene expression of genes from the ancestral linkage groups R, B1, B2, and C2, in three species: *P. flava* (left columns), *S. purpuratus* (middle columns), and *B. floridae* (right columns). Trees above the heatmaps represent the evolutionary history of each ALG in each deuterostome phylum. Trees below represent the similarity between species based on gower distance (calculated by similarity in broad timing of expression). Dots in most similar nodes indicate support for similarity calculated by the Adjusted Rand Index (fold change compared to the mean of all possible three pairwise comparisons > 1.5). Colour coding indicates broad timing of expression.

As shown in our companion manuscript, some of the Ancestral Linkage Groups (ALGs) conserved throughout bilateria underwent reorganisation in hemichordates and ambulacraria (Lin et al., in prep.). To explore changes in the expression of genes who underwent reshuffling during ambulacraria and hemichordate evolution, we explored the expression patterns of the genes belonging to ALGs R, B1, B2, and C2, in three species (*P. flava,* Sea Urchin, and *B. floridae*) (Fig. 6E, Sup. Fig. S17 E, Sup. Data 11). We observed the evolutionary history of each independent ALG is recapitulated by the gene expression of the genes in each species. For example, the R ALG underwent different changes in the three lineages: it fused and mixed with ALG B1 in hemichordates, it got spread into other chromosomes in chordates, and it remained unchanged in echinoderms. Although a number of genes kept their temporal order of gene expression, similarities in the gene expression of the genes in the three species was the lowest from these four ALGs. The B1 ALG underwent fusion and mixing only in hemichordates, and remained unchanged in echinoderms and chordates. Indeed we observed a higher similarity in the temporal order of genes between *B. floridae* and *S. purpuratus*, compared to the gaining of post-gastrulation expression in some genes of *P. flava*. The B2 and C2 ALGs fused and mixed with each other in the stem lineage that led to ambulacraria, while it remained unchanged in the stem lineage that led to chordates. In both ALGs, we found the highest similarities in the temporal order of genes between the two ambulacrarian species compared to *B. floridae*, with the two ambulacraria tending to share pre-gastrulation expression of genes otherwise expressed after gastrulation in amphioxus. These observations pose the question of the role of genomic rearrangement during the emergence of ambulacraria- or hemichordate-specific programmes of gene expression.

## Discussion

In this study we generated an extensive catalogue of transcriptomic and chromatin accessibility in the indirect-developing hemichordate *Ptychodera flava*. The morphologically distinct biphasic life cycle of *P. flava* is underlied by transcriptomic changes which also exhibit a biphasic behaviour, with gastrulation and the Muller (Tornaria) larva as the turning points of development. This transcriptional programme is likely controlled by two distinct TF programmes with little overlap as suggested by our coexpression and motif enrichment analyses.

During this biphasic programme, gastrulation seems to be the most evolutionarily conserved stage of hemichordate development, as indicated by the Transcriptomic Age Index and our comparative analyses. This could be related to the high interconnectivity of trans-developmental genes as inferred from our cis-regulatory network analyses, which might impose evolutionary constraints in this stage. These constraints might also be reflected in the conservation of gene expression with other deuterostome species such as cephalochordates and echinoderms. Similarities with other deuterostomes can potentially inform the equivalent timing of different processes during development in each deuterostome lineage. For example, similar gene family usage between ambulacraria larvae suggests a common indirect-developing ambulacrarian ancestor as they deploy similar expression programmes during these stages (B. J. Swalla 2006; E. Röttinger and Martindale 2011). Similarities with cephalochordates hint at related stages for the timing of neurogenesis and organogenesis. Conservations between hemichordate gastrula and amphioxus neurula hint at a similar mechanism for the establishment of the basis of the nervous system in the two lineages (Kaul-Strehlow and Röttinger 2015; Miyamoto and Wada 2013; Nielsen and Hay-Schmidt 2007). This does not contradict the idea that the hemichordate adult nervous system might originate from the rearrangement of components of the larval nervous system (Kaul-Strehlow et al. 2015). More data on *P. flava* and other deuterostomes, preferably at the tissue or cell type-level and spanning further into metamorphic development, are required to explore the significance of these observations in order to better identify and determine the conservation degree of gene expression, and to compare post-gastrulation and adult tissues with those of echinoderms and cephalochordates. These could potentially hint at the degree of gene module usage and the conservation of homologous structures and mechanisms, such as the aforementioned larval structures.

These similarities also extend to the expression of regulatory genes such as transcription factors. The presence of most of the cis-regulatory interactions between the genes of the endomesoderm kernel, supported by the conserved cis-regulatory activity of FoxA OCRs, supports the idea that the endomesoderm kernel emerged before the onset of ambulacraria. The presence of cis-regulatory interactions for the axial mesoderm regulatory network might indicate that the potential for these interactions pre-date the origin of chordates and were present in the last common ancestor of deuterostomes, although further functional studies are required to test this hypothesis. Of note is to mention that, since hemichordates lack a notochord homologous structure (Eric Röttinger and Lowe 2012; Lowe 2021), new questions arise as to which structure deploys the axial mesoderm GRN in hemichordates. Potential candidates include the endodermal region and protocoel, since some of the orthologous genes within the network are known to express in these embryonic territories. More validation is needed using *in situ* hybridisation to corroborate the co-expression or co-localization of genes belonging to the same GRNs. Likewise, the full scale of the GRNs has not been elucidated because our methods are based on cis-regulatory interactions and not protein-protein interactions (such as ligands and receptors), a known limitation of this analysis. The presence of broad temporal timing of HMG and bZIP classes before and after gastrulation, as observed between deuterostome species, might hint at the deuterostome LCA having a similar temporal programme of gene expression. Overall, these comparative analyses depict a more complete picture of the deuterostome LCA with the capacity for cis-regulatory interactions for neurulation, but also a biphasic life cycle controlled by an equally biphasic temporal trend of gene expression of TF classes, the emergence of which could also date back to the origin of bilateria if considering previous evidences from protostomes (Adryan and Teichmann 2010).

The agreement between gene expression and genomic closeness in the pharyngeal gene cluster of four deuterostome species is reassuring of the functional consolidation of this cluster at the onset of this lineage. The more divergent expression of the orthologues in the two protostome species, which we think is due to them not clustering together, suggests this co-regulation of gene expression in deuterostomes might be due to the presence of common cis-regulatory elements in the chromatin vicinity of the pharyngeal cluster genes. Overall, these analyses contribute to setting the precedent for future analysis of co-regulation and gene clustering to infer conservation between distant lineages.

Still, our analyses also support an independent evolutionary trajectory of hemichordate genome regulation, especially during larval development, and potentially underpinned by genomic rearrangements in the stem lineage of ambulacraria as well as the stem lineage of hemichordates. Post-gastrulation genes tend to be embedded in the cis-regulation that takes place during gastrulation to be deployed later in development, including TF genes that increase their centrality in the network and that belong to TF classes expressed later in development. These larval genes tend to be evolutionarily younger, as shown in our gene age enrichment and in the transcriptomic age index analyses. Since genes from the ALG that underwent independent mixing and fusion only in hemichordates tend to express later in development, we hypothesise that genomic rearrangement played a role in the evolution of larval strategies in hemichordates. More studies in other hemichordates, both direct- and indirect-developing species, will prove crucial to assess the degree and impact of these changes in the genome for the regulation during development.

In summary, this study presents the first high-throughput, genome wide, multimodal catalogue of quantitative regulatory genomics in hemichordates. In doing so, it establishes the basis for comparative studies in all deuterostome lineages with a global characterisation of the embryonic and larval development of an indirect-developing hemichordate species. Integration of this data with genome architecture analyses effectively generates hypotheses of the role of genome rearrangement in the evolution of hemichordate and deuterostome body plans.

## Materials and Methods

### Sample collection and animal husbandry

*Ptychodera* flava adults were collected from Penghu Islands, Taiwan. Methods for spawning induction and embryo culture were described previously (Lin et al. 2016). For RNA-seq, total RNA of each developmental stage was isolated from one to three biological replicates using the RNeasy Micro Kit (Qiagen). Library preparation for stranded polyA-based mRNA was then performed using the TruSeq Stranded mRNA Library Prep Kit (Illumina). The multiplexed cDNA libraries were subjected to sequencing with 151 bp paired-end read length on a HiSeq 2500 sequencer (Illumina). The average sequencing depth was 24,861,133 paired reads per sample. For ATAC-seq, embryos at different stages were collected to reach 75,000 cells per sample (∼700 early blastulae, ∼150 early gastrulae, ∼100 mid gastrulae and ∼50 late gastrulae). The collected embryos were treated with 0.1% N-acetylcysteine for 10 sec to remove the jelly coat, followed by incubating with 8.5 mM of DTT in seawater containing 5.7 mM of NaOH. The cells were then lysed for the transposition reaction and PCR amplification, and the purified libraries were sequenced as described (Marlétaz et al. 2018).

### Time-series analysis RNA-seq analysis

To quantify gene expression, reads were pseudo aligned against the set of longest transcripts per isoform per gene using Kallisto (Bray et al., n.d.). RNA-seq counts were corrected for batch effect and quantile normalised using edgeR (Robinson et al., n.d.; McCarthy, Chen, and Smyth 2012; Chen, Lun, and Smyth 2016) and RUVSeq (Risso et al. 2014) taking the 10000 bottom-ranked genes in edgeR as in-silico empirical, invariable genes. Pairwise Spearman correlation of replicates was done and visualised in R (Ripley 2001) using base and ComplexHeatmap (Gu, Eils, and Schlesner 2016) packages. Differential Gene Expression Analysis (DGEA) was performed using DESeq2 (Love, Huber, and Anders 2014) using likelihood-ratio-test and RUV-correction as reduced formula. Genes with an adjusted p-value below 0.05 (negative binomial LRT) were taken as differentially expressed. For every replicate, variance-stabilised counts were averaged to generate a time-series dataset of every differentially expressed gene throughout development. For every gene, expression data was scaled on a z-score (standard deviation over mean). Stage-specific clusters were then generated using hierarchical clustering on the scaled dataset. Data visualisation was done using R base (https://www.r-project.org/), ggplot2 (Wickham 2011), RColorBrewer (Neuwirth 2022), colorspace (Zeileis et al. 2019), pheatmap (RRID:SCR_016418), and ComplexHeatmap (Gu, Eils, and Schlesner 2016) packages under R 4.0.3.

### Orthology and Gene Age assignment

We ran OMA standalone (Altenhoff et al. 2021) using the set of predicted protein sequences of Ptychodera and a database of proteomes from 22 species (Sup. Data 4), using standard parameters. The resulting Orthogroups matrix was used to assign a gene age to each orthogroup (and therefore to every *Ptychodera* gene in a given orthogroup) based on Dollo parsimony using the Count software (Csurös 2010). We also parsed the .orthoxml output file of Hierarchical Orthogroups to assign a gene family ID to every *Ptychodera*, amphioxus, and sea urchin gene. These gene id - gene family ID association tables were used in our comparative analysis. As an alternative method for orthology inference, we ran OrthoFinder (Emms and Kelly 2015, 2019) using standard parameters on the same set of proteomes as we did for OMA. We retrieved one-to-one orthologs from the results, and also parsed the ‘Orthogroups.tsv’ output file to assign a gene family ID to every *P. flava*, amphioxus, and sea urchin gene. In addition, we also ran BLASTp best reciprocal hits between all species of interest (see “Comparative analysis” below, including *P. flava* and *B. floridae*) to retrieve one-to-one orthologues.

### Gene age Enrichment analysis and Transcriptomic Age Index analysis

We performed gene age enrichment analysis using contingency tables for every cluster and gene age. Results were considered significant when Fisher test p-value was lower than 0.01. We performed transcriptome age index analysis using the R package ‘myTAÌ and tested for significance using the flat-line test.

### ATAC-seq analysis

ATAC-seq data analyses were performed as described in (Gil-Gálvez et al. 2022), using standard pipelines (Marlétaz et al. 2018; Buenrostro et al. 2013). Briefly, reads were aligned with Bowtie2 using a de novo sequenced, HiC-guided assembly of the genome of *Ptychodera flava* (deposited; see Data Availability). We generated a dataset of peaks using Macs2 (Y. Zhang et al. 2008) for peak calling and Bedtools (Quinlan and Hall 2010) to quantify the accessibility of each chromatin region. Differential analysis was performed using DESeq2 v1.18.0 in R 3.4.3 (Love, Huber, and Anders 2014) using likelihood-ratio test (LRT). An adjusted p-value = 0.05 was set as a cutoff for statistical significance of the differential accessibility of the chromatin in ATAC-seq peaks. Motif enrichment was calculated using the programme ‘FindMotifsGenome.pl‘ from the Homer tool suite (Heinz et al. 2010). k-means clustering of the ATAC-seq signal was performed using R (Ripley 2001) and visualised using ComplexHeatmap (Gu, Eils, and Schlesner 2016). Open Chromatin Regions were assigned to the closest transcription start site (TSS) using bedtools closestbed and standard parameters. Categorisation of OCRs into distal, proximal, intronic etc., as well as *de novo* motif finding and motif enrichment analysis, was done using the program ‘annotatePeaks‘ from the Homer tool suite (Heinz et al. 2010).

### Cis-regulatory reporter assays

The ATAC-seq data revealed 4 potential REs locating within 3 kb upstream of the *foxa* coding sequence. PCR products containing 4, 3, 2, or 1 REs were cloned into the Hugo’s lamprey construct (HLC) vector (Parker, Bronner, and Krumlauf 2014; Minor et al. 2019) upstream of an *egfp* coding sequence and an SV40 polyadenylation sequence using In-Fusion Snap Assembly master mixes (Takara). To prepare the injection mixtures, 4 ng of reporter constructs (PCR products), 1.2 ul of 1M KCl, 154 ng of carrier DNA (HindIII digested *S. purpuratus* gDNA), and 5 ug of Dextran Alexa 555 were added into nuclease-free water (Invitrogen) to make 10 ul of mixture. About 4 pl of this mix was introduced into *S. purpuratus* zygotes, and the EGFP signal was observed at blastula and gastrula stages.

### Gene annotation

#### Generation of Protein sequences

We ran TransDecoder (https://bio.tools/TransDecoder) using the set of longest isoforms per gene of Ptychodera, querying against the SWISSPROT and Pfam databases to gather evidence at the sequence homology and protein domain level. This evidence was subsequently incorporated into Transdecoder and we retrieved the longest predicted protein sequences with the parameter --single-best-only.

#### Gene Ontology Annotation

Using the TransDecoder-predicted set of protein sequences, GO terms were generated using a combination of BLAST2GO and eggNOG annotation software, using RefSeq non-redundant proteins (NR) Database (Pruitt, Tatusova, and Maglott 2007) and eggNOG metazoa (Huerta-Cepas et al. 2019) as reference databases respectively.

#### Gene Ontology (GO) enrichment analysis

For every cluster, GO enrichment analysis was performed using the topGO R package (Alexa, Rahnenführer, and Lengauer 2006; “topGO” n.d.) using the whole gene set of *Ptychodera* as gene universe and the *elim* method (Alexa, Rahnenführer, and Lengauer 2006).

#### Trans-developmental and housekeeping classification

A gene was classified as trans-developmental if annotated with any of the following GO terms: GO:0044430, GO:0006412, GO:0003735, GO:0005840, GO:0005842, GO:0030529, GO:0006414, GO:0009119, GO:0009062, GO:0044262, GO:0006725; but none of the GO terms for housekeeping genes. A gene was classified as housekeeping if annotated with any of the following GO terms: GO:0009790, GO:0030154, GO:0043565, GO:0007267, GO:0008380, GO:0003700, GO:0140297, GO:0010467, GO:0010468, GO:0016070; but none of the GO terms for trans-developmental genes.

### Transcription Factor annotation

We first generated groups of orthology using *Ptychodera*, Purple Sea Urchin, Japanese Feather Star, amphioxus, and well-annotated vertebrate species (human, zebrafish) using OrthoFinder (Emms and Kelly 2015, 2019) (Sup. Data 4). Using human and zebrafish queries for TFDB Metazoa (Hui Hu et al. 2019), we transferred TF class annotation to their respective orthogroup and thus to every gene of the other species. Secondly, we also added to this classification all genes identified as a TF homolog by the GimmeMotifs tool ‘motif2factors‘ (briefly, transferring TF annotation based on orthology evidence using OrthoFinder against a standard set of model species; see gimmemotifs documentation), which was required for ANANSE (see methods below). The Resulting TF annotation was used for downstream TF analysis.

### Transcription Factor analysis

For a given TF class X in a given species (*Ptychodera*, Amphioxus, Sea Urchin) at a given developmental stage Y, we computed i) the sum of genes of TF class X expressed (> 1 TPM) at stage Y; ii) the sum of TPMs (using kallisto (Bray et al. 2016), see above) for every expressed gene of class X at stage Y; and iii) the rate of ii divided by i. To measure correlation between classes and stage-specific gene expression, we first computed the average expression profile (z-scored) of every stage-specific cluster. Then, for a given TF class X, we computed the number of TFs of class X with a similar expression profile (z-scored) as that of the stage specific cluster (with a Spearman correlation above 0.6), divided by the number of TFs in class X. We did this for every TF class with every stage-specific cluster. For motif discovery analysis, we first extracted the open chromatin regions (differential or not) associated with genes of each stage-specific cluster of gene expression (pre-larval development). Then, we performed motif enrichment analysis using the programme FindMotifsGenome.pl from the Homer tool suite (Heinz et al. 2010).

### Cis-regulatory networks

We computed two networks of cis-regulation (early blastula and late gastrula) using ANANSE (Xu et al. 2021). We initially generated a custom database of TF-domain association using GimmeMotifs ‘motif2factors‘ (Bruse and van Heeringen 2018), tailored for the genome annotation of *Ptychodera*. For each developmental stage, we first ran ANANSE binding to create a genome-wide profile of TF binding across regions of open chromatin using the .bam alignment of the ATAC-seq, and then ran ANANSE network to create a network of transcription factors (TFs) and target genes (TGs) using the binding profile and the RNA-seq expression data. The resulting network was imported to R and analysed using iGraph. We subsetted the network to retrieve edges of the top decile (ANANSE score value, ‘prob’, above quantile 0.9), and subsequently computed network size, number of connections per TG, and per-TFclass centrality using custom wrappers for iGraph functions in R. Germ layer genes were retrieved from in situ hybridisation data on hemichordates in the scientific literature, and assigned to the newest annotation using BLAST best reciprocal hits. Genes from the notochord / paraxial mesoderm GRNs were retrieved from the scientific literature and used as query on a BLAST best-reciprocal hit against the newest annotation of *Ptychodera*. Code and full documentation is available at the online repository of the project.

#### *In situ* hybridisations

*In situ* hybridisations were performed using embryos of *Ptychodera flava* at the late gastrula stage, as previously described in (Ikuta et al. 2013; Luo and Su 2012; Lu, Luo, and Yu 2012).

### Reanalysis of Amphioxus and Sea Urchin

#### Reanalysis of Sea Urchin

We retrieved RNA-Seq reads of *Strongylocentrotus purpuratus* from (Tu, Cameron, and Davidson 2014) and mapped them against the Spur5.0 (Arshinoff et al. 2022) set of longest isoforms per gene using kallisto (Bray et al. 2016). Read counts were imported to R using the ‘tximport‘ package. Counts were transformed to account for batch effect using edgeR and RUVseq as explained above, and we used edgeR to identify the top variable genes in lieu of performing differential gene expression analysis due to lack of replicates in the original dataset (the only sample with two replicates, Mesenchyme Blastula, was kept as separate replicates for all the analyses). We clustered these highly-variable genes in R using hierarchical clustering. We also annotated the TF repertoire of Sea Urchin and measured TF class composition of each transcriptome as well as measured correlation between classes and stage-specific gene expression, as explained above for *Ptychodera*. Code and full documentation is available at the online repository of the project.

#### Reanalysis of Branchiostoma lanceolatum

RNA-Seq reads of *Branchiostoma lanceolatum* from (Marlétaz et al. 2018) were mapped against the set of longest isoforms per gene of the currently available transcriptome (Brasó-Vives et al. 2022) using kallisto (Bray et al. 2016). Read counts were imported to R using the ‘tximport‘ package. We used ‘DESeq2‘ to identify temporally expressed genes using the negative binomial likelihood ratio test as explained above. We clustered the resulting significant genes in R using hierarchical clustering to retrieve clusters of stage-specific genes. We also annotated the TF repertoire of *B. lanceolatum* and measured TF class composition of each transcriptome as well as measured correlation between classes and stage-specific gene expression, as explained above for *Ptychodera*. Code and full documentation is available at the online repository of the project.

#### Reanalysis of Anneissia japonica

RNA-Seq reads of *Anneissia japonica* from (Li et al. 2020) were mapped against the set of longest isoforms per gene of the currently available transcriptome using kallisto (Bray et al. 2016). Read counts were imported to R using the ‘tximport‘ package. We used ‘DESeq2‘ to identify temporally expressed genes using the negative binomial likelihood ratio test as explained above. We clustered the resulting significant genes in R using hierarchical clustering to retrieve clusters of stage-specific genes. Code and full documentation is available at the online repository of the project.

#### Reanalysis of Branchiostoma floridae

RNA-Seq reads of *Branchiostoma floridae* from (Haiyang Hu et al. 2017) were mapped against the set of longest isoforms per gene of the currently available transcriptome using kallisto (Bray et al. 2016). Read counts were imported to R using the ‘tximport‘ package. We used ‘DESeq2‘ to identify temporally expressed genes using the negative binomial likelihood ratio test as explained above. We clustered the resulting significant genes in R using hierarchical clustering to retrieve clusters of stage-specific genes. Code and full documentation is available at the online repository of the project.

### Comparative analyses

Using our OMA output, we retrieved all one-to-one orthologues between *P. flava* and *S. purpuratus*, between *P. flava* and *A. japonica*, and between *P. flava* and *B. lanceolatum*. For a given pair of species, we used a set of custom wrappers we called ‘comparABlè (https://github.com/apposada/comparABle), that incorporate well-established methods in the literature (such as pairwise transcriptomic comparisons and orthogroup overlap; see below), to perform comparative analyses between these pairs of species. Alternatively, we also used the Orthofinder and BLASTp reciprocal best hits outputs to retrieve one-to-one orthologues between these species and run the comparative analysis. Code and full documentation is available at the online repository of the project.

#### Pairwise transcriptomic correlations

Using the whole temporal transcriptome datasets of each species, we subsetted these datasets to retrieve only common one-to-one orthologs and calculated distance and correlation metrics (Pearson correlation, Spearman correlation, and Jensen-Shannon Divergence) between pairs of transcriptomic stages. We calculated the average Jensen-Shannon Divergence using bootstrapping of 1000 iterations with replacement, and also performed co-occurrence analysis (Levy et al. 2021; Álvarez-Campos et al. 2023) using all genes and bootstrapping with replacement using the concatenate of the two datasets.

#### Identification of genes driving similarities between stages

Using the most closely matched pairs of stages according to the correlations described above, as well as selecting particularly interesting stage pairs, we identified key genes expressed during specific stage combinations across various species. Briefly, for a pair of stages a.i and b.j, we generated a design matrix to compare a.i and b.j against the rest of stages, performed linear modelling on the concatenate dataset of one-to one orthologues using limma (Ritchie et al. 2015), and retrieved the top genes with p.value <0.05. Because this method finds genes significantly expressed in these two stages and not the others, it effectively retrieves a subset of the genes that drive similarities between stages a.i and b.j in a conservative manner, at the expense of other genes conspicuously expressed which might also be important for these similarities even if they are not captured by this method.

#### Gene family usage analysis

We compared all possible combinations of clusters of stage-specific genes between pairs of species by performing orthogroup overlap analysis (Martín-Zamora et al. 2023; Marlétaz et al. 2018). For a given cluster in species ‘a’ a.x and a given cluster in species ‘b’ b.y, we counted the number of overlapped orthogroups between a.x and b.y (the number of genes from a given orthogroup in clusters a.x and b.y), and subjected these to upper-tail hypergeometric test. Using the information of the pairs of clusters sharing the highest amount of orthogroups, we retrieved the genes from Ptychodera belonging to those orthogroups and performed functional category analysis using the Clusters of Orthologous Groups annotation derived from EggNOG (see methods above).

#### Comparative analyses using transcription factors

We generated reciprocal best hits of P. flava vs B. lanceolatum and P. flava vs. S. purpuratus using BLASTp with standard parameters, and used these as proxy for one-to-one orthologs as an alternative method to OMA or OrthoFinder. Upon seeing similar trends regardless of using OMA or reciprocal best hits, we used reciprocal best hits to identify conserved TFs between the two pairs of species. We subsetted the expression datasets to keep the one-to-one TFs and performed the same distance metric correlations between pairs of species. Using the information on the most similar stages we proceeded with linear modelling to retrieve the TF genes driving similarities between stages as explained above.

#### Transcriptomic data of pharyngeal-related genes

Time-series transcriptomic data from eight species based on previous studies were downloaded from NCBI (Sup. Data 2), including for amphioxus (PFL), sea urchin (SPU), sea star (POC and PMI), oyster (CGI), scallop (PYE), jellyfish (CHE) and sponge (AQU). Illumina adaptors and low-quality bases were removed using Trimmomatic (version 0.36) (Bolger, Lohse, and Usadel 2014). The trimmed reads were then mapped to the corresponding reference sequences using STAR aligner (version 2.7.10a). The alignment files combined with the gene model files were subjected to raw counts estimation using featureCounts software (version 2.0.1) (Liao, Smyth, and Shi 2014). Expression levels were normalised as TPM (Transcripts Per Million). Expression levels of pharyngeal-related genes were then extracted and subjected to visualisation using MeV software (version 4.9.0) (Howe et al. 2011).

#### Comparison of gene expression in ALG groups

We retrieved the *P.flava*, *S. purpuratus* and *B. floridae* orthologues of genes found in the ALGs of the last common ancestor of deuterostomes (see Methods from (Lin et al., in prep.)). For every of these genes, we assigned a broad timing of expression based on their classification in one of each clusters of stage-specific gene expression of *P. flava*, *B. floridae*, and *S. purpuratus*. For the latter, we expanded the assignment of genes to clusters by correlating the expression of every gene in the S. purpuratus transcriptome against all the average expression profiles of the stage-specific clusters of S. purpuratus. Genes with a max correlation of 0.7 or higher to a given cluster were assigned to that cluster. We plotted the expression of these genes using the package ComplexHeatmap. We calculated species similarity using Gower distance (using the daisy function from the R package cluster) in the matrix of genes x per-species behavior matrices, with genes in rows and per-species behaviour (timing before or after gastrulation) in columns. Adjusted Rand Index between methods of grouping genes by broad timing in pairs of species (e.g. agreement between the timing of expression between *P. flava* orthologues and *B. floridae* orthologues) was calculated using the adjustedRandIndex from the R package mclust.

## Supplementary Data

All Supplementary Data information is publicly available at: https://figshare.com/projects/Ptychodera_supplementary_data/141296.

All code and figures are publicly available at: https://github.com/apposada/ptychodera_cisreg_development

## Data availability

P. flava genome assembly used in this work is publicly available: https://www.ncbi.nlm.nih.gov/bioproject/PRJNA747109. The version described in this paper is version JASXRY010000000.

Raw and processed sequencing data were deposited in GEO (GSE207835, token for reviewer: ypslgsccldkfvex).

## Acknowledgements

We thank Yi-Cheng Chang, Yi-Chih Chen, Chang-Tai Tsai and Cindy Chou for collecting *P. flava* samples for RNA-seq and ATAC-seq. We thank the NGS High Throughput Genomics Core, Biodiversity Research Center, Academia Sinica. This study was supported by the Ministry of Science and Technology, Taiwan (Grant MOST-110-2326-B-001-006 to Y.H.S. and Grant MOST-110-2621-B-001-001-MY3 to J.K.Y.) and Academia Sinica (AS-GC-111-L01 to Y.H.S. and J.K.Y.). J.J.T. and J.L.G.S. were supported by the European Research Council (ERC, grant agreement No 740041) and the Spanish Ministerio de Economía y Competitividad (grant PID2019-103921GB-I00). We thank Alejandro Gil Galvez and Panos Firbas for advice with the analyses, and Maarten Van der Sande, Siebren Frolich, and Simon Van Heeringen, for their advice with ANANSE. A.P.P. especially thanks the rest of the authors and all the members of the Skarmeta/Tena lab, as well as Jordi Solana, for their trust and support during the difficult times of this project.

## Work by Authors

A.P.P. performed the RNA-seq, ATAC-seq, TF analyses, Network analyses, orthology analyses, comparative analyses, and ALG expression analyses, and wrote the first version of the manuscript. Che Y.L. generated the genome annotation and generated the ALG data (see companion manuscript). Ching Y.L. and Y.C.C. performed de *In situ* hybridisations. T.P.F. performed the cis-regulatory reporter assays. J.L.G.S., Y.H.S. and J.K.Y. conceptualised the project. J.J.T., Y.H.S. and J.K.Y. supervised and provided feedback on the analysis and the writing. All authors edited the manuscript.

## Notes

https://figshare.com/projects/Ptychodera_supplementary_data/141296

### Competing Interest Statement

The authors have declared no competing interest.

### Summary of Updates

The manuscript has been rewritten, with new datasets incorporated into the analyses. Figures have been reorganized, and two additional figures with further analyses have been included.

