## Supplementary Figures for "Insights into deuterostome evolution from the biphasic transcriptional programme of hemichordates"

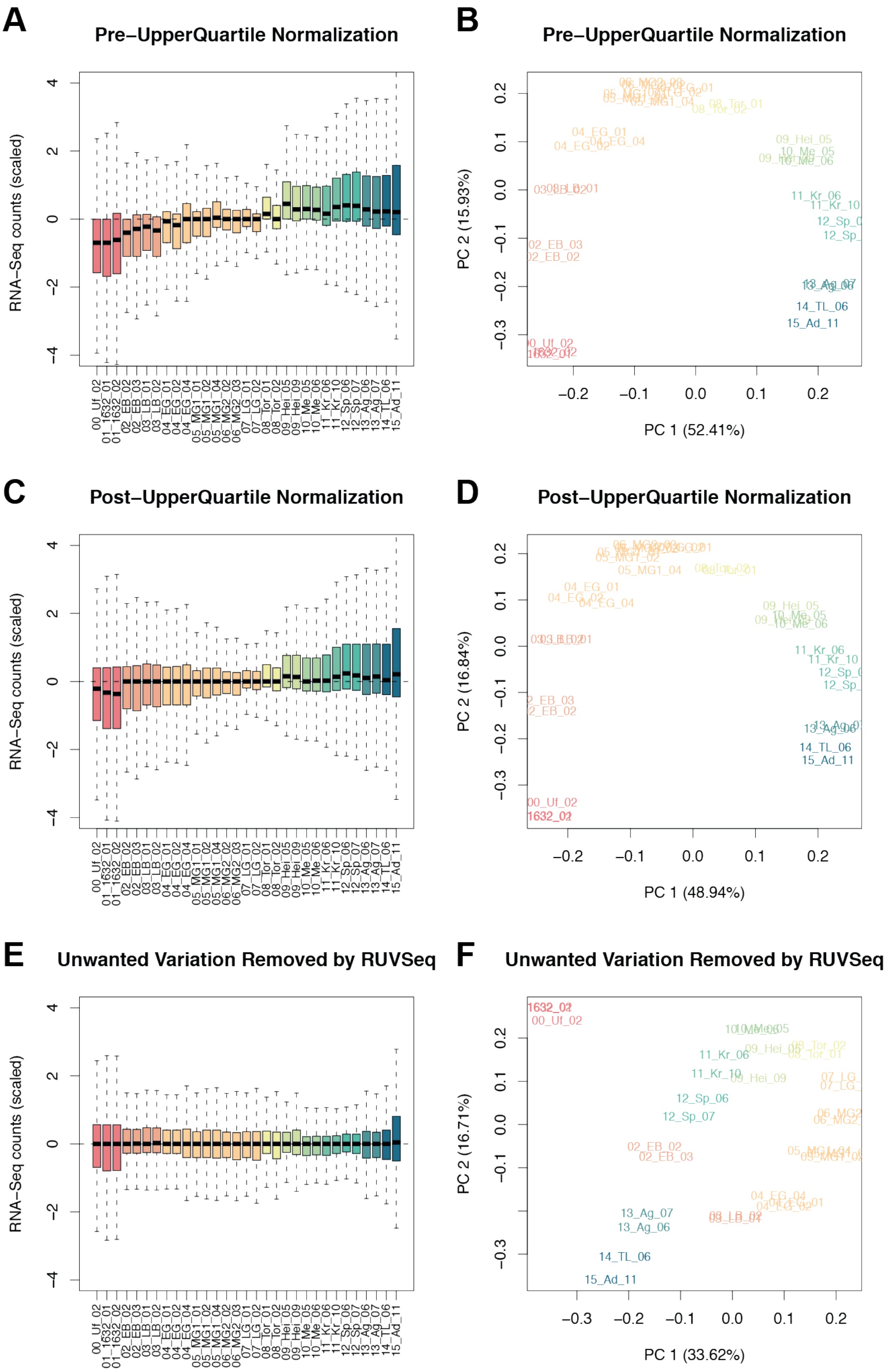


**Supplementary Figure S1.** Count distribution (A,C,D) and principal component analysis (B,D,F) for the developmental transcriptomes before upper quartile normalisation (A,B), after upper quartile normalisation (C,D), and after applying RUVSeq (E,F).

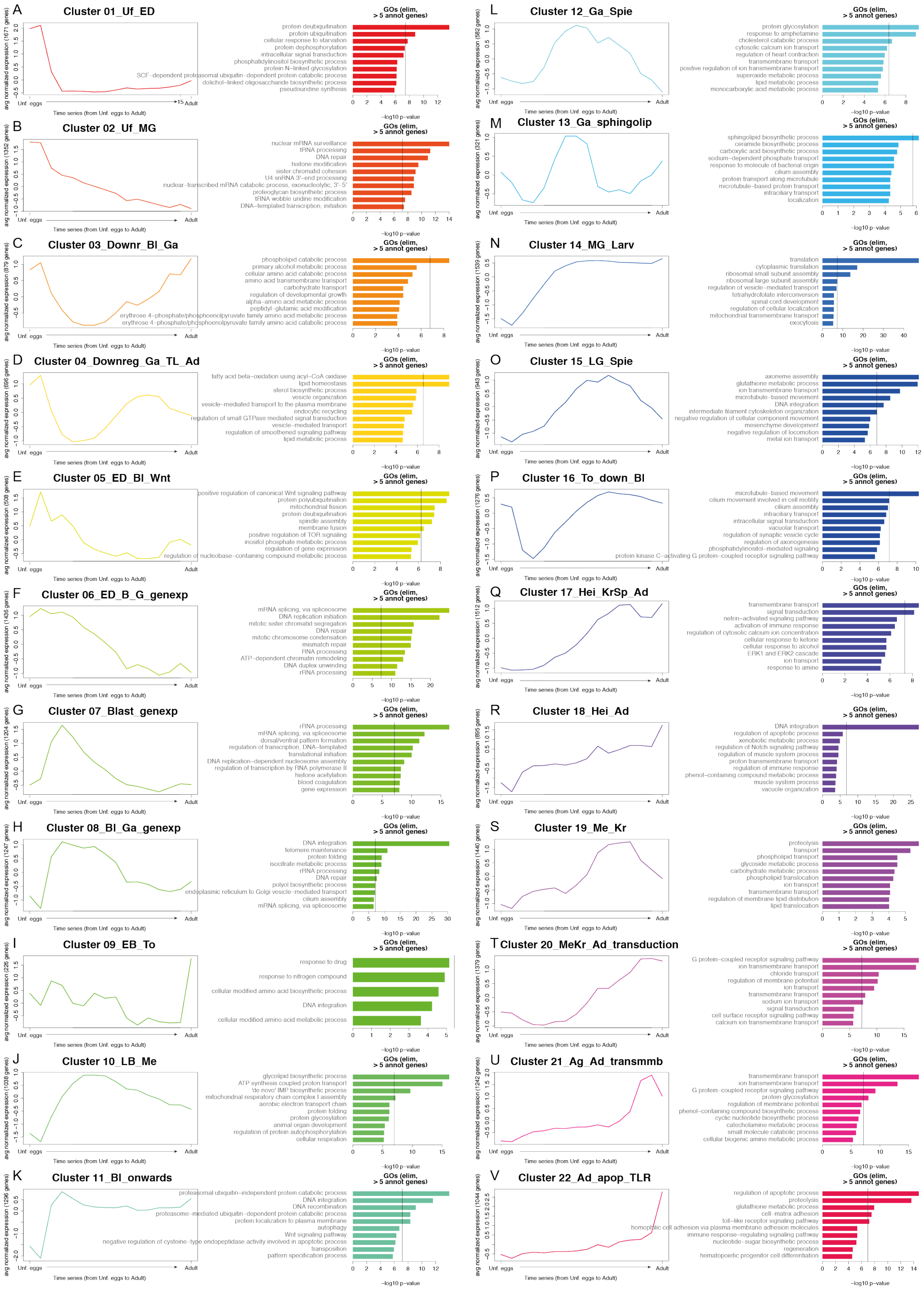

**Supplementary Figure S2.** For each cluster of stage-specific genes (A-V), Left: averaged expression profile of the genes in each cluster, and Right: Barplot of top enriched GO terms.

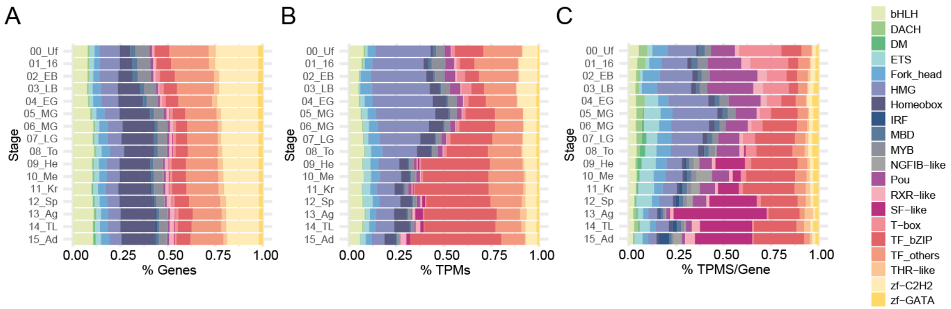


**Supplementary Figure S3.** Stacked barplot showing the TF composition of each developmental stage of Ptychodera in terms of number of TFs from each class (A), fraction TPMs per TF class (B), and fraction of TPMs per gene per class (C).

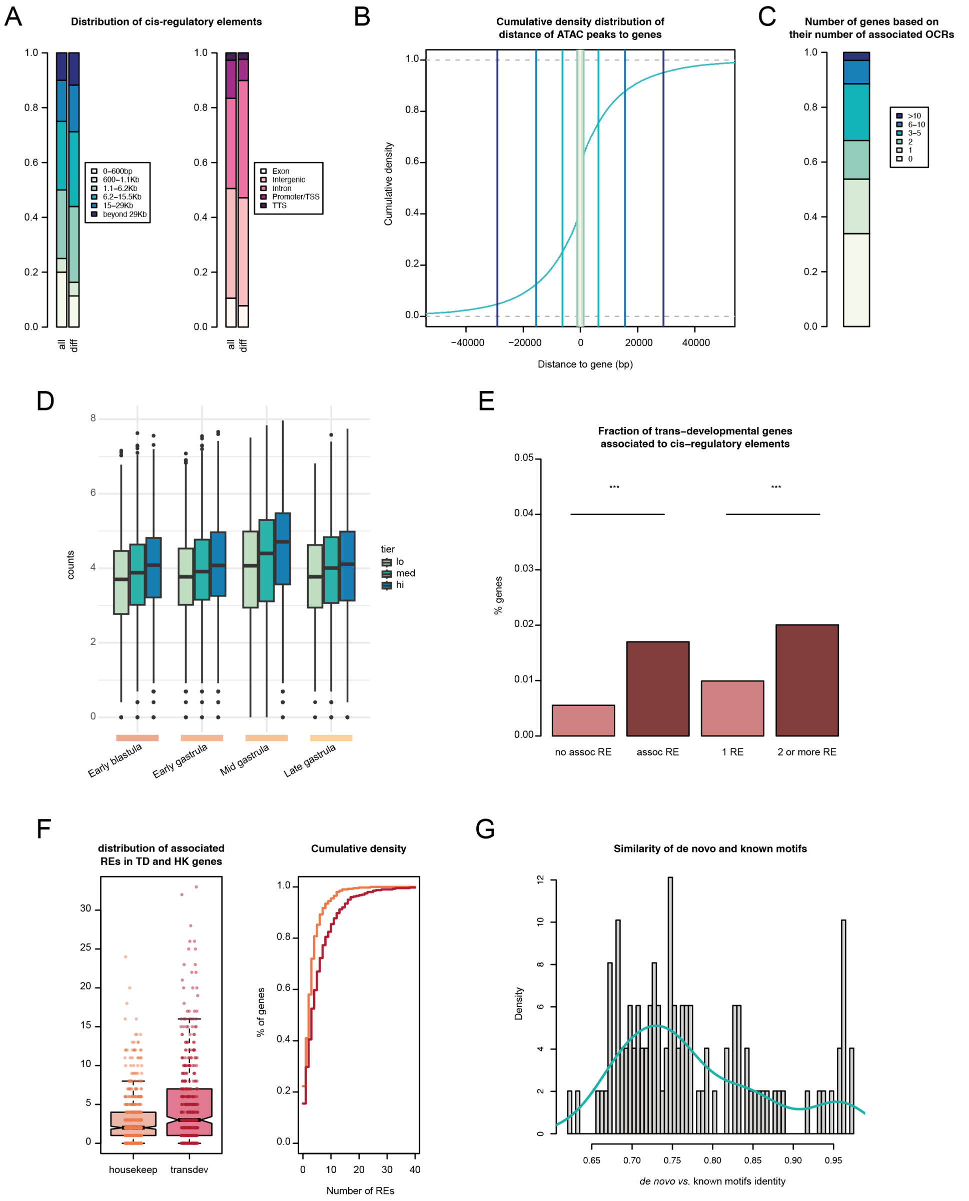


**Supplementary Figure S4.** Characterisation of ATAC-seq dataset of Ptychodera flava during gastrulation. A: Distribution of open chromatin regions by relative distance to the TSS (left), and by categorisation in different genomic features (right). B: Cumulative density curve of open chromatin regions relative to the transcription start site of genes. Blue lines indicate quantile thresholds used for classification in panel A. C: Distribution of genes in relation to the number of associated open chromatin regions. D: Boxplots of counts in open chromatin regions of each developmental stage, divided in tiers regarding the level of expression of their associated genes (lo=low expression, med=medium expression, hi=high expression). E: Fraction of trans-developmental genes (see Methods) associated to open chromatin regions. F: Left: boxplot of the number of open chromatin regions associated to housekeeping or trans-developmental genes (N= 400). Right: Cumulative density function of number of genes in relation to the number of associated OCRs. x axis: number of associated OCRs; y axis: fraction of genes in the population. G: Histogram and density plot of the similarity scores of two hundred de-novo inferred motifs in the genome of Ptychodera, in relation to known vertebrate motifs (from the databases included in the HOMER software).

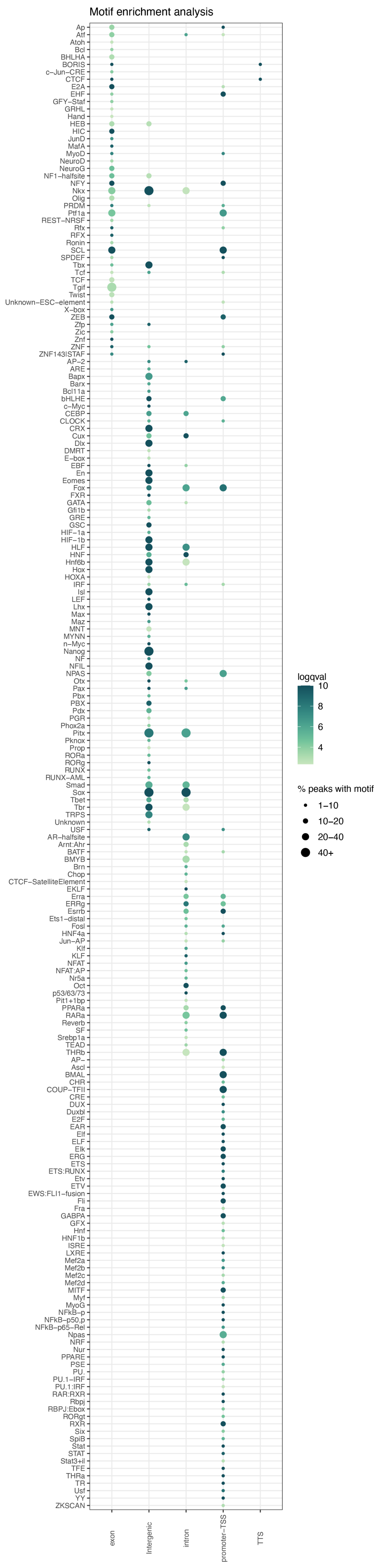


**Supplementary Figure S5.** Box plot depicting motif enrichment analysis in the open chromatin regions classified as different genomic features (exon, intergenic, intron, promoter/TSS area, and TTS. Dot size represents fraction of OCRs with motif found, and colour indicates log-qvalue (hypergeometric test).

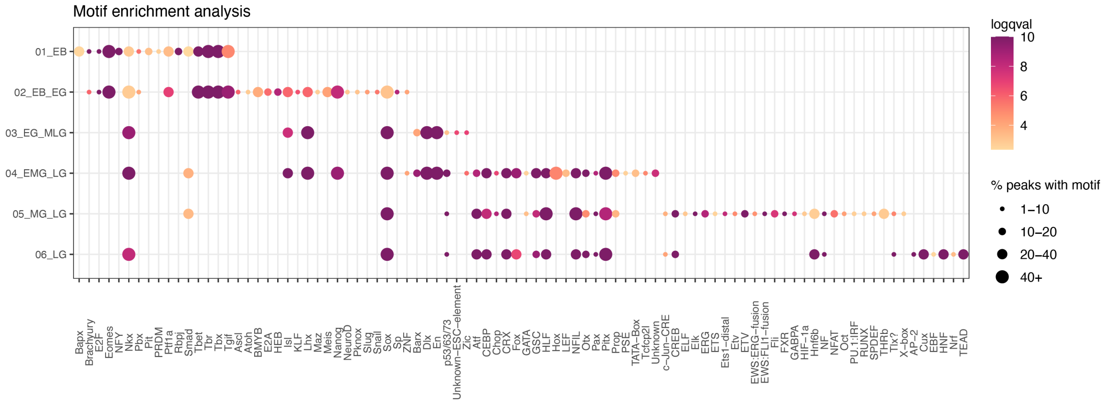


**Supplementary Figure S6.** Box plot depicting motif enrichment analysis in differentially accessible regions of open chromatin. Dot size represents fraction of OCRs with motif found, and colour indicates log-qvalue (hypergeometric test).

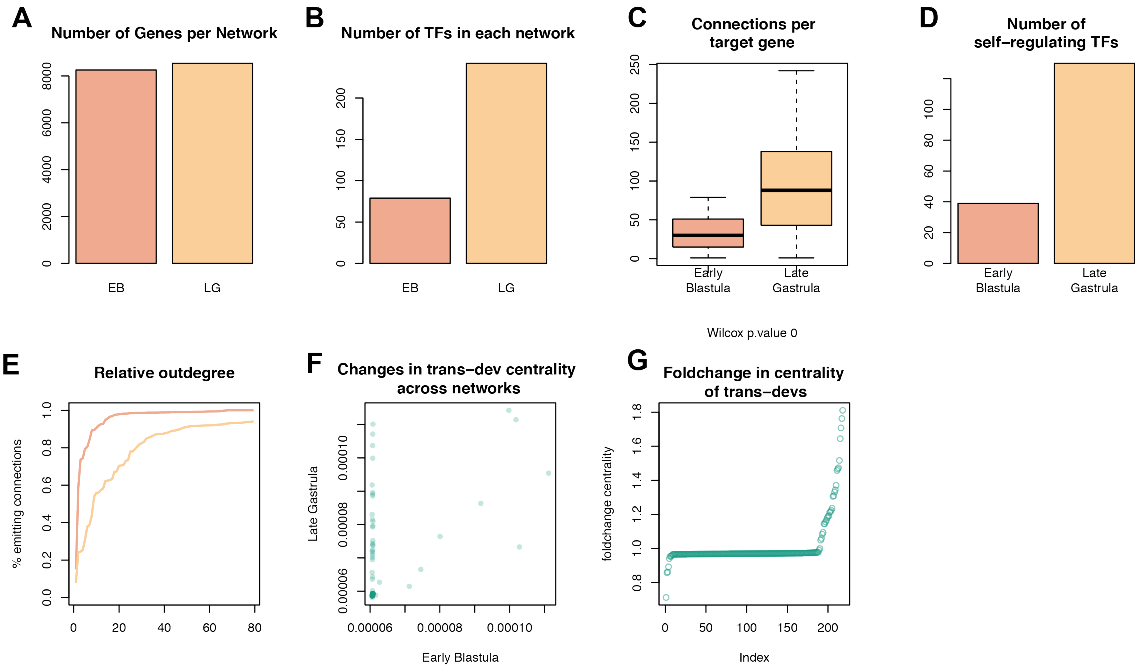


**Supplementary Figure S7.** Network metrics of Ptychodera gastrulation. A: Number of genes per network. B: Number of Transcription factors embedded in each network. C: Boxplot showing the distibution of edges (connections) per target gene in each network. D: Number of self-regulating TFs (TF genes with a binding motif of itself in its proximity). E: Cumulative distributions of the relative outdegree of TFs (fraction of emitted connections of a given gene over the total connections involving that gene) in each stage. F: Scatter plot of the centrality of trans-developmental genes in each stage. X axis: centrality in early blastula; Y axis: centrality in late gastrula. G: fold-change of trans-dev centrality in. X axis: sorting index, Y axis: fold-change of centrality. Each data point is a gene.

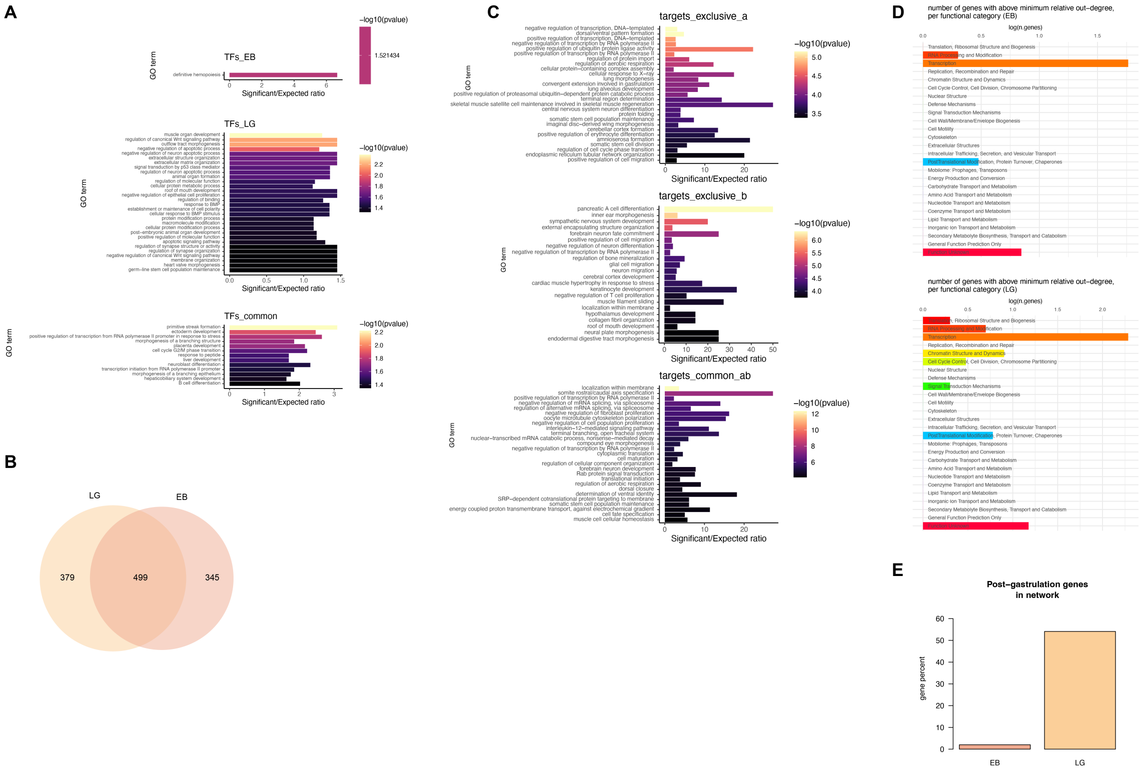


**Supplementary Figure S8.** A: barplots of GO term enrichment for early blastula-specific, late gastrula-specific, and common TF genes in the networks. Bar size represents foldchange of observed/expected genes and colour intensity represents -log10(qvalue). B: Venn diagram showing the number of common and specific target genes of each network. C: barplots of GO term enrichment for early blastula-specific, late gastrula-specific, and common target genes in the networks. Bar size represents foldchange of observed/expected genes and colour intensity represents -log10(qvalue). D: Barplot showing the number of genes of each functional category that show an above-minimum relative outdegree in each of the networks. X axis: functional categories (COG); y axis: log of number of genes. E: Percentage of genes from post-gastrulation clusters, present in each network. Y axis: gene percentage (out of the total of genes classified as post-gastrulation stages-specific clusters). F: sub-graph of influential TF genes as calculated by ANANSE influence. Node size represents level of expression, edge thickness represents interaction score.

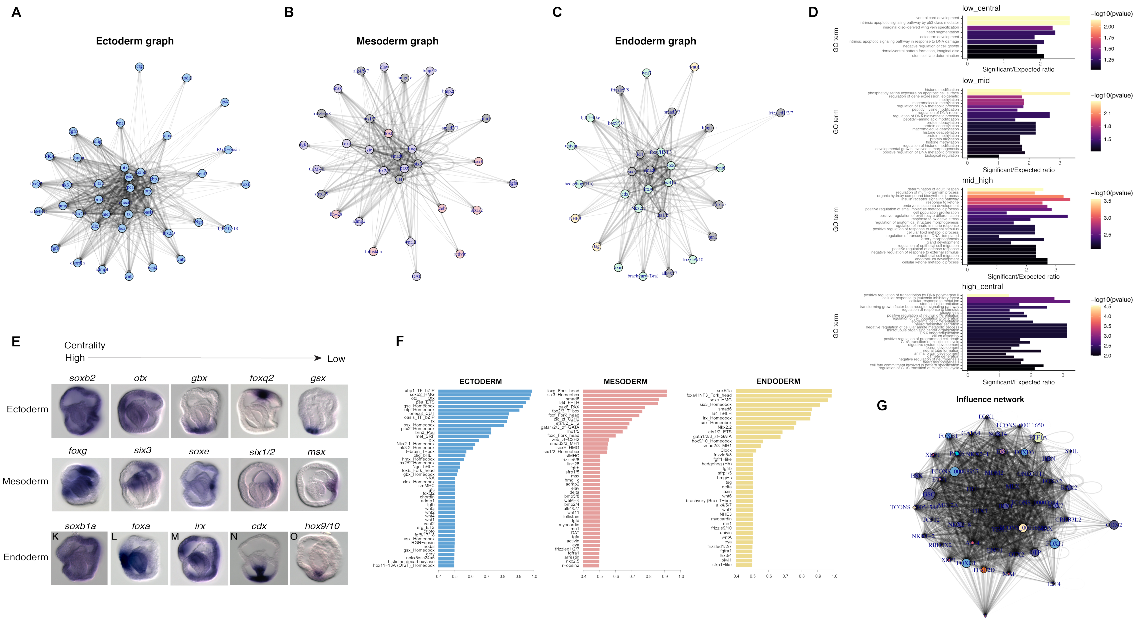


**Supplementary Figure S9.** A,B,C: Sub-graphs of genes found to be expressed in ectoderm (A), mesoderm (B), and endoderm (C) based on in situ hybridisation experiments. D: centrality values for different transcription factors in the network, based on the germ layer where they are expressed. E: barplots of GO term enrichment for low centrality, low-to-mid centrality, mid-to-high centrality, and high centrality TF genes in the networks. Bar size represents foldchange of observed/expected genes and colour intensity represents -log10(qvalue). F: *In situ* hybridisations of TF genes expressed in different germ layers, showing an agreement (inverse relationship) between embryonic territory specificity and centrality in the network.

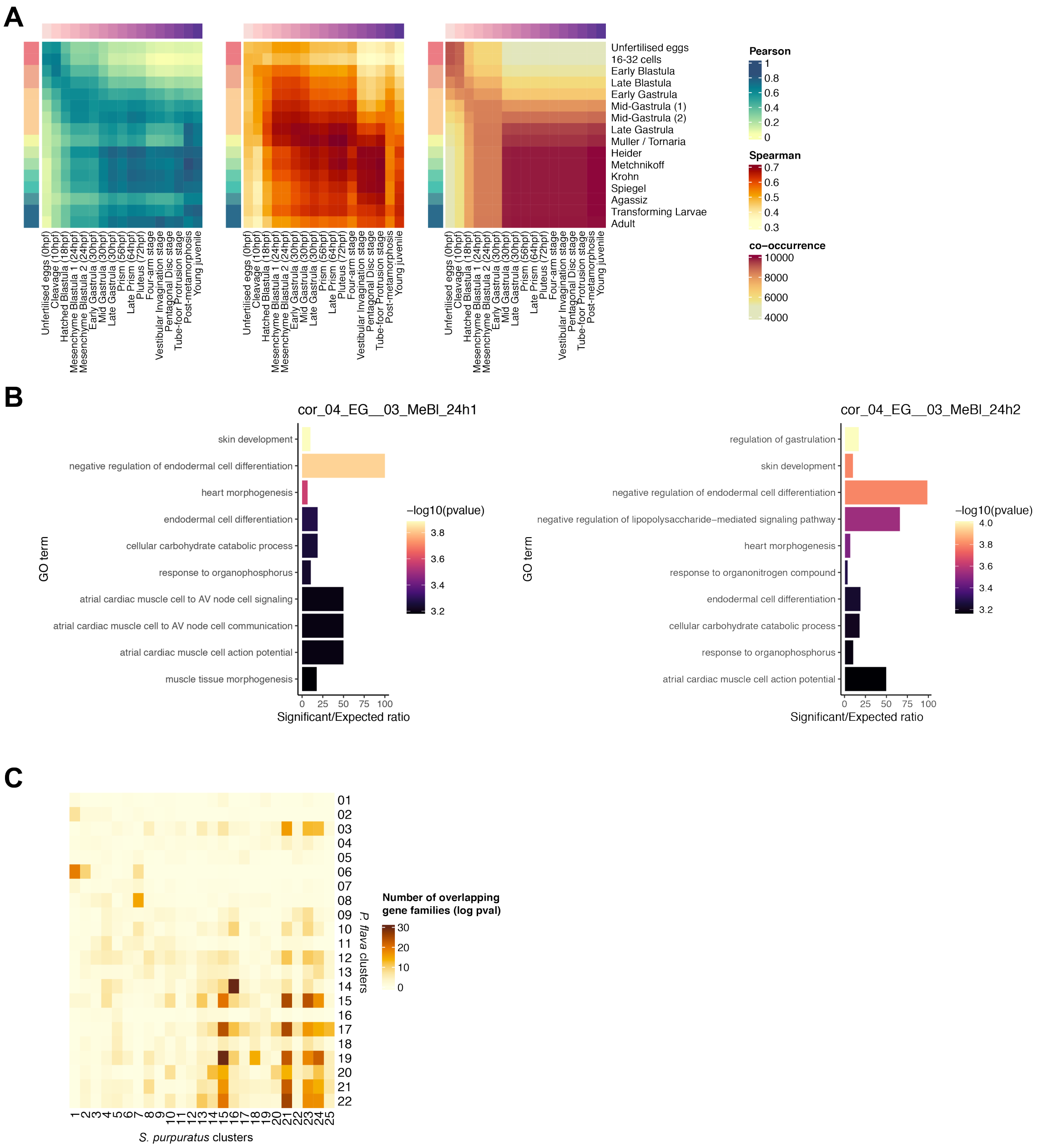


**Supplementary Figure S10.** A: Pearson correlation (left), Spearman correlation (right), and co-occurrency (right) of pairwise comparisons of developmental transcriptomes between Ptychodera flava (rows) and Purple sea urchin (columns). B: Gene Ontology terms of genes found specifically expressed in the stages of hemichordate Early Gastrula and Mesenchyme Blastula. C: orthology overlap in pairwise comparisons of gene families from stage-specific clusters. Colour intensity indicates -logpvalue of the upper tail hypergeometric test.

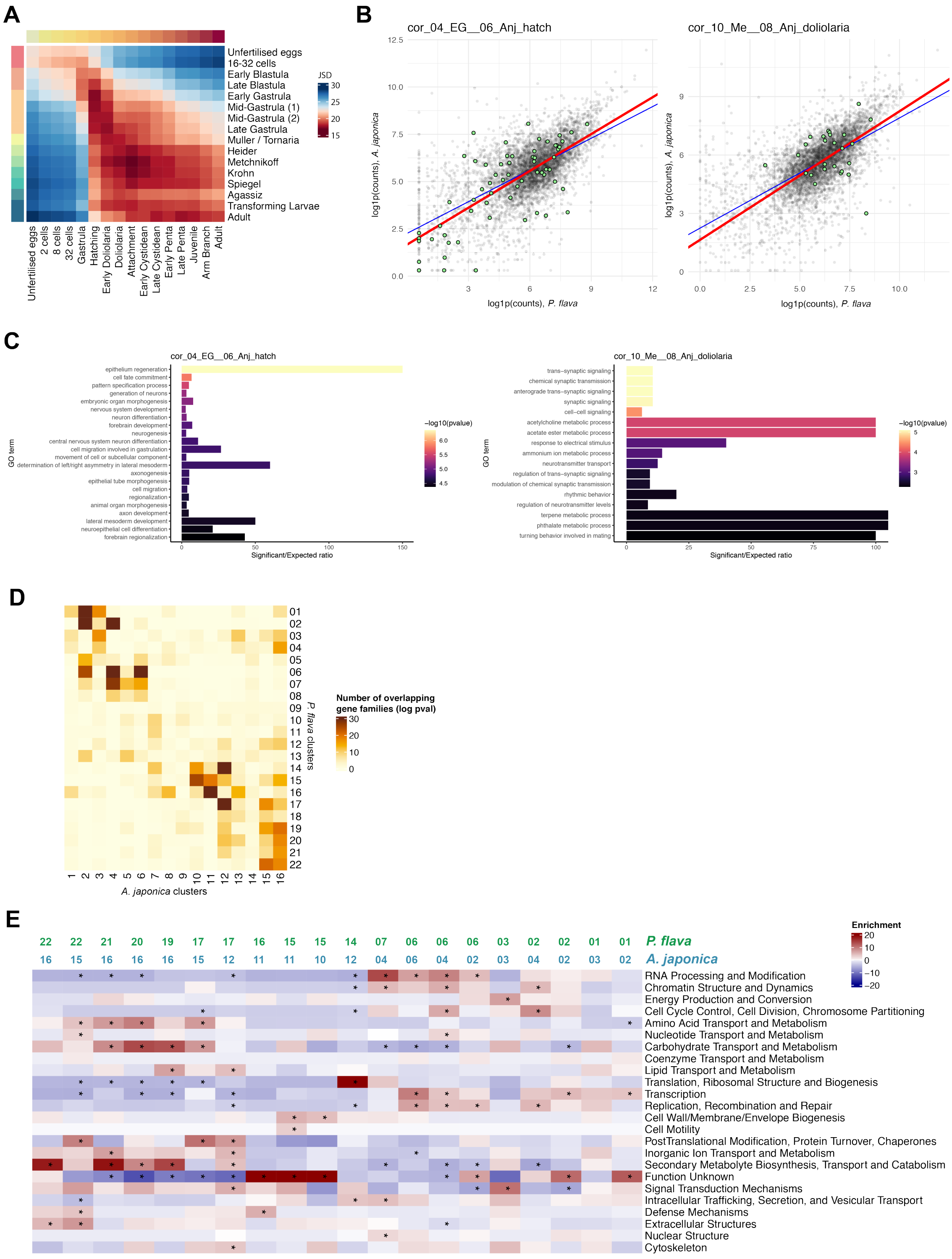


**Supplementary Figure S11.** Comparative transcriptomics of *Ptychodera flava* and the crinoid *Anneissia japonica*. A: Jensen-Shannon divergence between transcriptomic stages of *P. flava* (rows) and *Anneissia japonica* (columns). B: scatter plot showing genes found commonly and exclusively expressed between hemichordate gastrula and crinoid hatching (left), and between hemichordate Metchnikoff larva and crinoid doliolaria (right). Green highlights the genes found expressed in these pairs of stages and not others; i.e. the “unique genes" responsible for the similarities observed (see Methods) C: Gene Ontology terms of genes found specifically expressed in the same pairwise comparisons as in B. D: orthology overlap in pairwise comparisons of gene families from stage-specific clusters. Colour intensity indicates -logpvalue of the upper tail hypergeometric test. E: heatmap showing functional category enrichment in gene families found common between different pairs of stage-specific clusters. Colour indicates percentage of enrichment. Asterisks indicate Fisher's test <0.05.

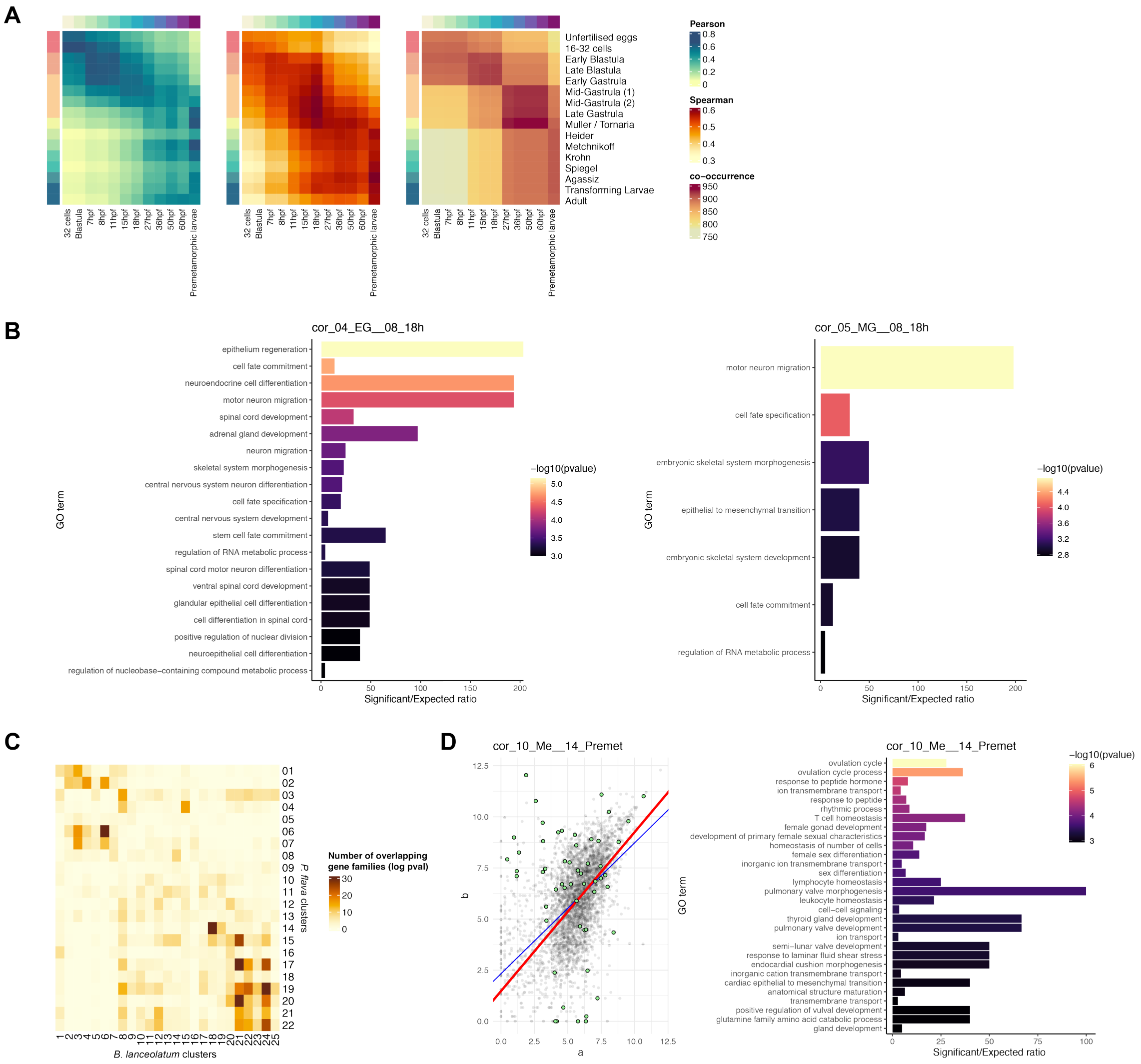


**Supplementary Figure S12.** A: Pearson correlation (left), Spearman correlation (right), and co-occurrency (right) of pairwise comparisons of developmental transcriptomes between Ptychodera flava (rows) and Amphioxus (columns). B: Gene Ontology terms of genes found specifically expressed in the stages of hemichordate Early Gastrula and 18hpf. C: orthology overlap in pairwise comparisons of gene families from stage-specific clusters. Colour intensity indicates -logpvalue of the upper tail hypergeometric test. D: same as A,B but between Metchnikoff larva from Ptychodera and the Pre-metamorphic larva of Amphioxus.

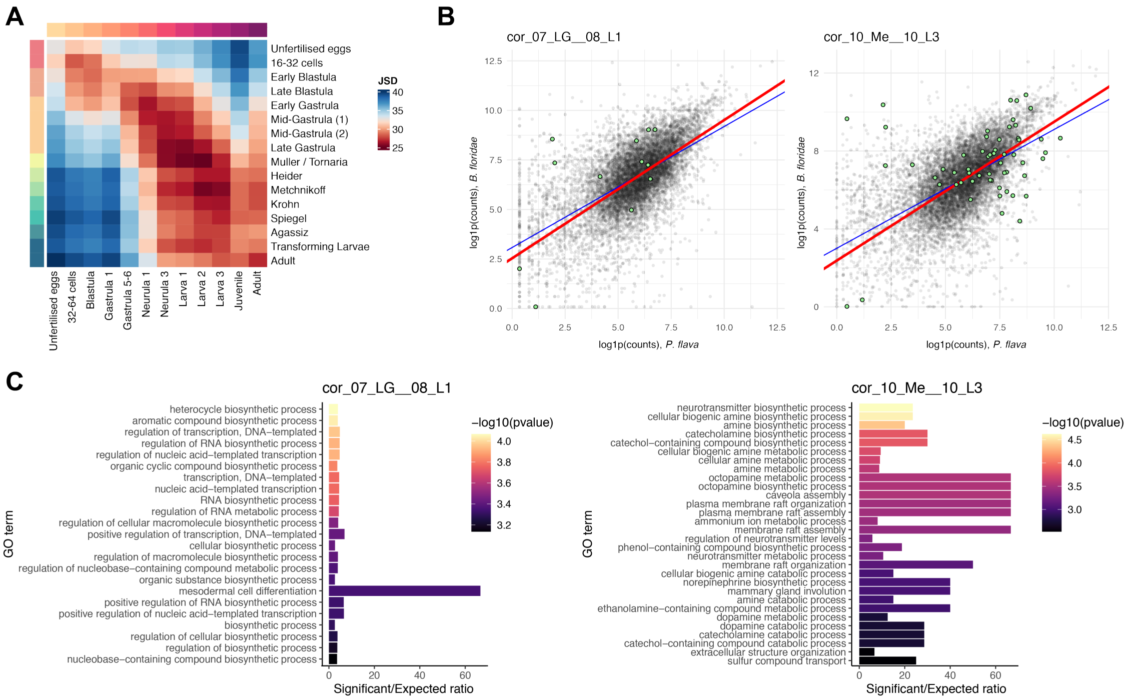


**Supplementary Figure S13.** Comparative transcriptomics of *Ptychodera flava* and the lancelet *Branchiostoma floridae*. A: Jensen-Shannon divergence between transcriptomic stages of *P. flava* (rows) and *B. floridae* (columns). B: scatter plot showing genes found commonly and exclusively expressed between hemichordate late gastrula and *B. floridae* Larva 1 (left), and between hemichordate Metchnikoff larva and *B. floridae* Larva 3 (right). Green highlights the genes found expressed in these pairs of stages and not others; i.e. the “unique genes” responsible for the similarities observed (see Methods). C: Gene Ontology terms of genes found specifically expressed in the same pairwise comparisons as in B.

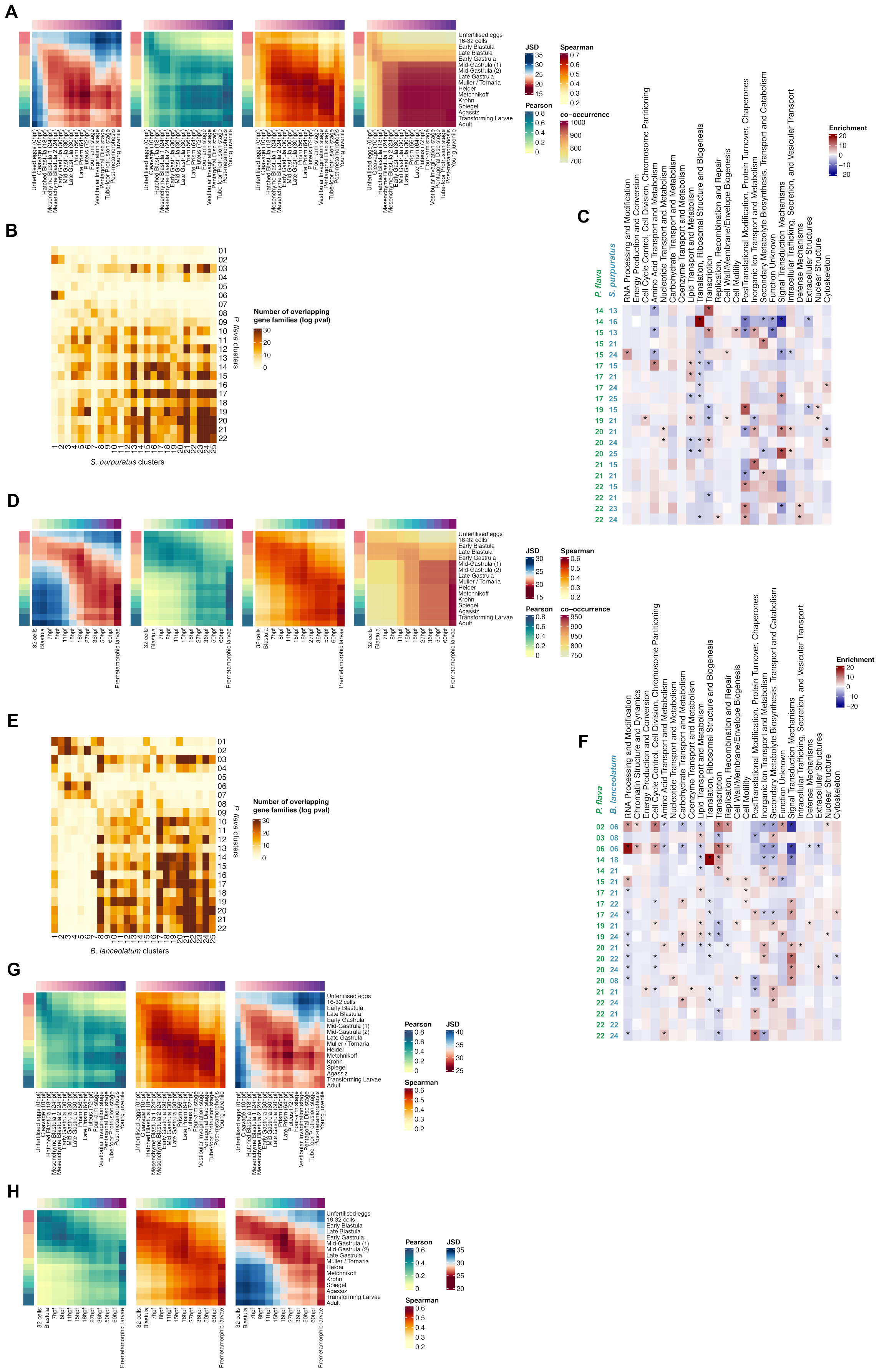
A
JSD
Spearman
Pearson
co−occurrence
Unfertilised eggs
16-32 cells
Early Blastula
Late Blastula
Early Gastrula
Mid-Gastrula (1)
Mid-Gastrula (2)
Late Gastrula
Muller / Tornaria
Heider
Metchnikoff
Krohn
Spiegel
Agassiz
Transforming Larvae
Adult
B
Number of overlapping gene families (log pval)
P. flava clusters
S. purpuratus clusters
C
P. flava
S. purpuratus
Enrichment
RNA Processing and Modification
Energy Production and Conversion
Cell Cycle Control, Cell Division, Chromosome Partitioning
Amino Acid Transport and Metabolism
Nucleotide Transport and Metabolism
Carbohydrate Transport and Metabolism
Coenzyme Transport and Metabolism
Lipid Transport and Metabolism
Translation, Ribosomal Structure and Biogenesis
Transcription
Replication, Recombination and Repair
Cell Wall/Membrane/Envelope Biogenesis
Cell Motility
PostTranslational Modification, Protein Turnover, Chaperones
Inorganic Ion Transport and Metabolism
Secondary Metabolyte Biosynthesis, Transport and Catabolism
Function Unknown
Signal Transduction Mechanisms
Intracellular Trafficking, Secretion, and Vesicular Transport
Defense Mechanisms
Extracellular Structures
Nuclear Structure
Cytoskeleton
D
32 cells
Blastula
Premetamorphic larvae
E
B. lanceolatum clusters
F
B. lanceolatum
Chromatin Structure and Dynamics
G
H

**Supplementary Figure S14.** (A,B,C) Reanalysis of *Ptychodera flava* and Sea Urchin comparative transcriptomics using Orthofinder results. A: from left to right: Jensen-Shannon Divergence, Pearson correlation, Spearman correlation, and Co-occurrence, of *P. flava* stages (rows) and Sea urchin stages (columns). B: orthology overlap in pairwise comparisons of gene families from stage-specific clusters. Colour intensity indicates -logpvalue of the upper tail hypergeometric test. C: heatmap showing functional category enrichment in gene families found common between different pairs of stage-specific clusters. Colour indicates percentage of enrichment. Asterisks indicate Fisher's test <0.05. (D,E,F) Reanalysis of Ptychodera and B. lanceolatum comparative transcriptomics using Orthofinder results. D: from left to right: Jensen-Shannon Divergence, Pearson correlation, Spearman correlation, and Co-occurrence, of Ptychodera stages (rows) and Sea urchin stages (columns). E: orthology overlap in pairwise comparisons of gene families from stage-specific clusters. Colour intensity indicates -logpvalue of the upper tail hypergeometric test. F: heatmap showing functional category enrichment in gene families found common between different pairs of stage-specific clusters. Colour indicates percentage of enrichment. Asterisks indicate Fisher's test <0.05. (G,H) Comparative transcriptomics between Ptychodera and Sea Urchin (G) or between Ptychodera and Amphioxus (H) using BLAST reciprocal best hits.

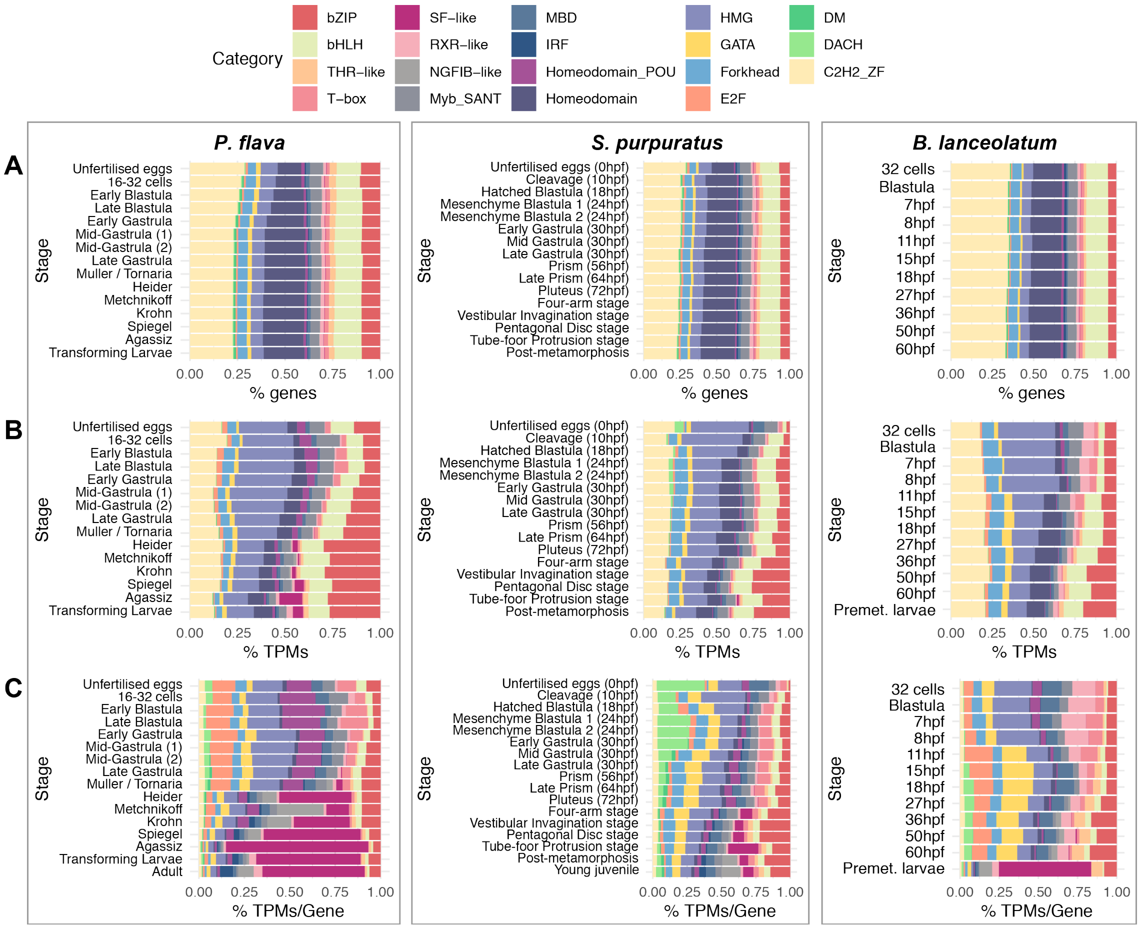


**Supplementary Figure S15.** A: Stacked barplot showing the TF composition of each developmental stage of Ptychodera (left), sea urchin (middle) and amphioxus (right) in terms of number of TFs from each class (A), fraction TPMs per TF class (B), and fraction of TPMs per gene per class (C). Ptychodera data are the same shown in Sup. Fig. S3.

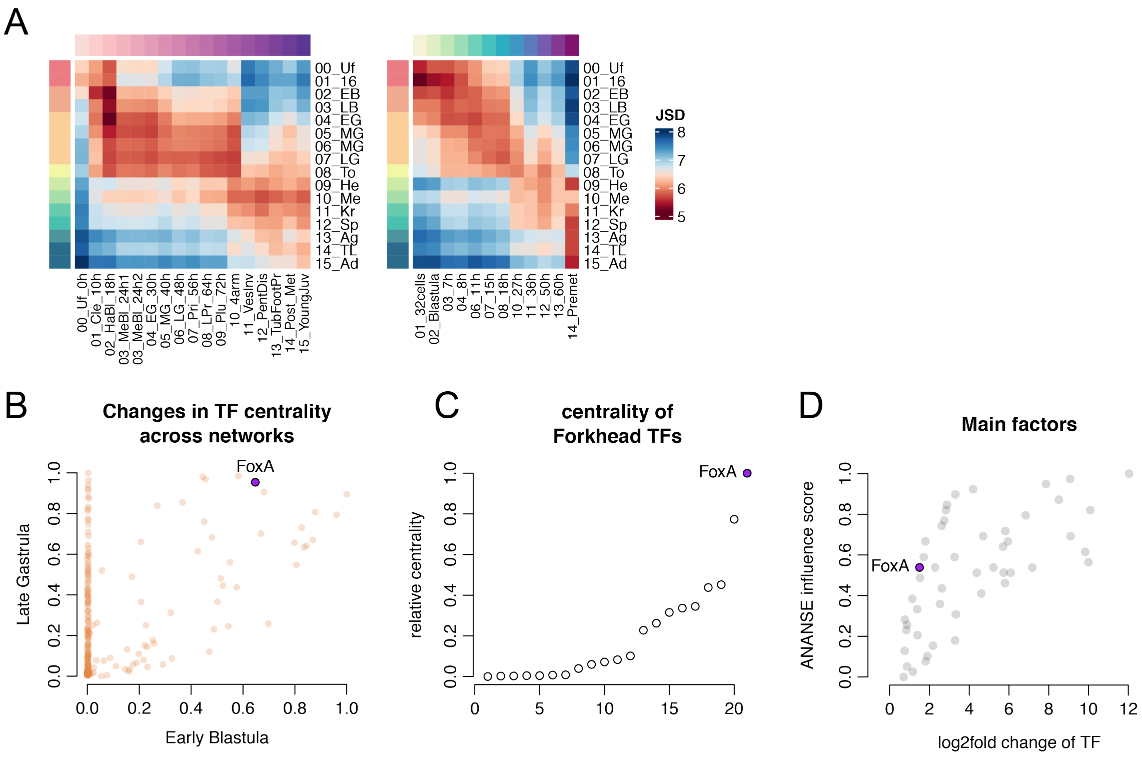


**Supplementary Figure S16.** A: Jensen Shannon Divergence of transcriptomic stages between Ptychodera and sea urchin (left) and Ptychodera and B. lanceolatum (right) using TF orthologues. B: Scatter plot of TF centrality in each network indicating the position of FoxA. C: Relative centrality in the LG network of all the Forkhead TFs, with FoxA as the most central TF. D: Scatter plot showing the top influential TFs as calculated by ANANSE influence, showing the position of FoxA.

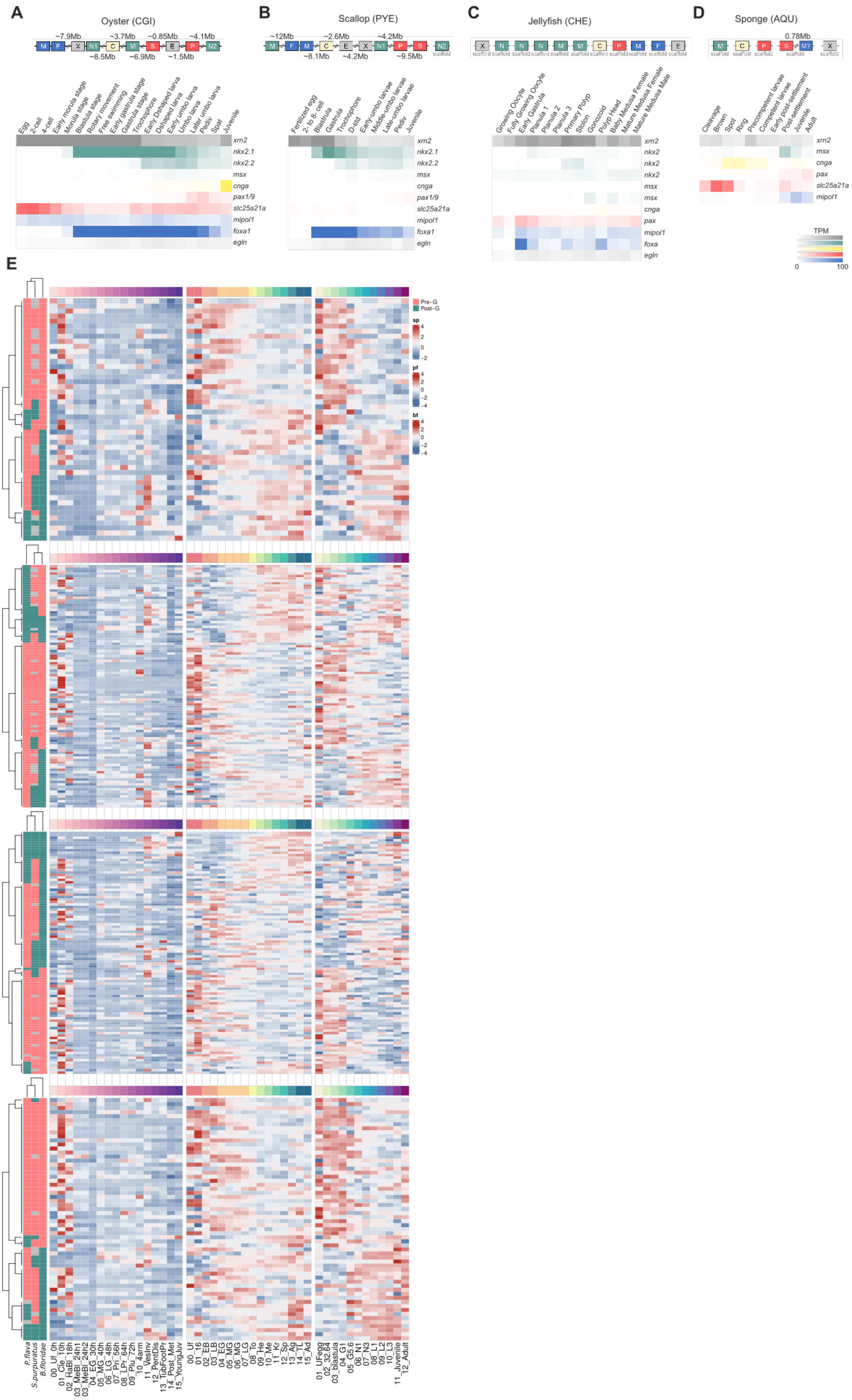

A
Oyster (CGI)
B
Scallop (PYE)
C
Jellyfish (CHE)
D
Sponge (AQU)
E
xrn2
nkx2.1
nkx2.2
msx
cnga
pax1/9
slc25a21a
mipol1
foxa1
egln
TPM
0
100
Pre-G
Post-G
P.flava
S.purpuratus
B.floridae

**Supplementary Figure S17.** A-D: heatmaps of gene expression of the pharyngeal gene cluster in protostome and early-branching metazoan species. E: heatmaps of gene expression of the genes from each ALG that underwent changes in deuterostome evolution, for the three species studied.
